# The PqsE active site as a target for small molecule antimicrobial agents against *Pseudomonas aeruginosa*

**DOI:** 10.1101/2022.04.21.489002

**Authors:** Isabelle R. Taylor, Philip D. Jeffrey, Dina A. Moustafa, Joanna B. Goldberg, Bonnie L. Bassler

**Affiliations:** Department of Molecular Biology, Princeton University, Princeton, NJ 08544, USA; Emory University School of Medicine, Children’s Healthcare of Atlanta, Inc., Department of Pediatrics, and Center for Cystic Fibrosis and Airway Diseases Research, Atlanta, GA 30322, USA; Howard Hughes Medical Institute, Chevy Chase, MD 20815, USA

**Keywords:** *Pseudomonas aeruginosa*, quorum sensing, antibiotics, protein-protein interactions, allosteric inhibitors

## Abstract

The opportunistic pathogen *Pseudomonas aeruginosa* causes antibiotic resistant, nosocomial infections in immuno-compromised individuals, and is a high priority for antimicrobial development. Key to pathogenicity in *P. aeruginosa* are biofilm formation and virulence factor production. Both traits are controlled by the cell-to-cell communication process called quorum sensing (QS). QS involves the synthesis, release, and population-wide detection of signal molecules called autoinducers. We previously reported that activity of the RhlR QS transcription factor depends on a protein-protein interaction with the hydrolase, PqsE, and PqsE catalytic activity is dispensable for this interaction. Nonetheless, the PqsE-RhlR interaction could be disrupted by substitution of an active site glutamate residue with tryptophan (PqsE(E182W)). Here, we show that disruption of the PqsE-RhlR interaction via either the E182W change or alteration of PqsE surface residues that are essential for the interaction with RhlR, attenuates *P. aeruginosa* infection in a murine host. We use crystallography to characterize the conformational changes induced by the PqsE(E182W) substitution to define the mechanism underlying disruption of the PqsE-RhlR interaction. A loop rearrangement that repositions the E280 residue in PqsE(E182W) is responsible for the loss of interaction. We verify the implications garnered from the PqsE(E182W) structure using mutagenic, biochemical, and additional structural analyses. We present the next generation of molecules targeting the PqsE active site, including a structure of the tightest binding of these compounds, BB584, in complex with PqsE. The findings presented here provide insight for drug discovery against *P. aeruginosa* with PqsE as the target.

**Author Summary:** The human pathogen *Pseudomonas aeruginosa* is resistant to many currently used antibiotics, making it a burden of urgent clinical importance. *P. aeruginosa* pathogenicity is controlled by the bacterial cell-to-cell communication process called quorum sensing (QS). The function of one protein that controls *P. aeruginosa* QS-directed virulence, RhlR, requires a protein-protein interaction with an enzyme called PqsE. When PqsE is blocked from interacting with RhlR, *P. aeruginosa* is avirulent and incapable of infecting an animal host. Here, we validate the PqsE-RhlR interaction as a target for antibiotic development, and we present a mechanism for how such antibiotics could disrupt the PqsE-RhlR interaction. Discovery of new antibiotics would fulfill an unmet healthcare need by providing treatments to combat *P. aeruginosa* infections.

## Introduction

*Pseudomonas aeruginosa* is an opportunistic human pathogen that is responsible for untreatable infections in vulnerable, immuno-compromised individuals (1,2). As a member of the “ESKAPE” multi-drug resistant pathogens, *P. aeruginosa* is a high-priority target for the development of next generation antibiotics that function by new mechanisms of action (3,4). Features contributing to *P. aeruginosa* pathogenicity include the ability to produce virulence factors and form biofilms (5). These traits are controlled by the bacterial cell-cell communication process called quorum sensing (QS) (6,7).

QS involves the production, release, and population-wide detection and response to signal molecules called autoinducers (8). *P. aeruginosa* QS relies on two acyl-homoserine lactone (HSL) autoinducer production/detection systems, called Las and Rhl, and the Pqs alkyl quinolone autoinducer production/detection system (9–11). The Las system consists of the LasI autoinducer synthase and LasR receptor/transcription factor which, respectively, produce and detect the autoinducer 3-oxo-C12-HSL (12). The Rhl system consists of the RhlI autoinducer synthase and the RhlR receptor/transcription factor, which, respectively, produce and detect the autoinducer C4-HSL (13) (Figure 1a). The Pqs system consists of the PQS autoinducer, synthesized by PqsABCD, and its partner receptor/transcription factor, PqsR (14,15). The Las system sits at the top of the *P. aeruginosa* QS hierarchy and controls transcription of the components of the Rhl and Pqs systems (16). Curiously, RhlR function additionally depends on the metallo-*β*-hydrolase enzyme PqsE, encoded as the final gene in the *pqsABCDE* biosynthetic operon (17–19). With respect to RhlR, PqsE is required to form a protein-protein interaction that enhances the affinity of RhlR for target promoters (20,21). Moreover, the RhlR-PqsE interaction is essential for *P. aeruginosa* virulence phenotypes, and therefore, of interest as a focus for antibiotic development.

**Figure 1:**
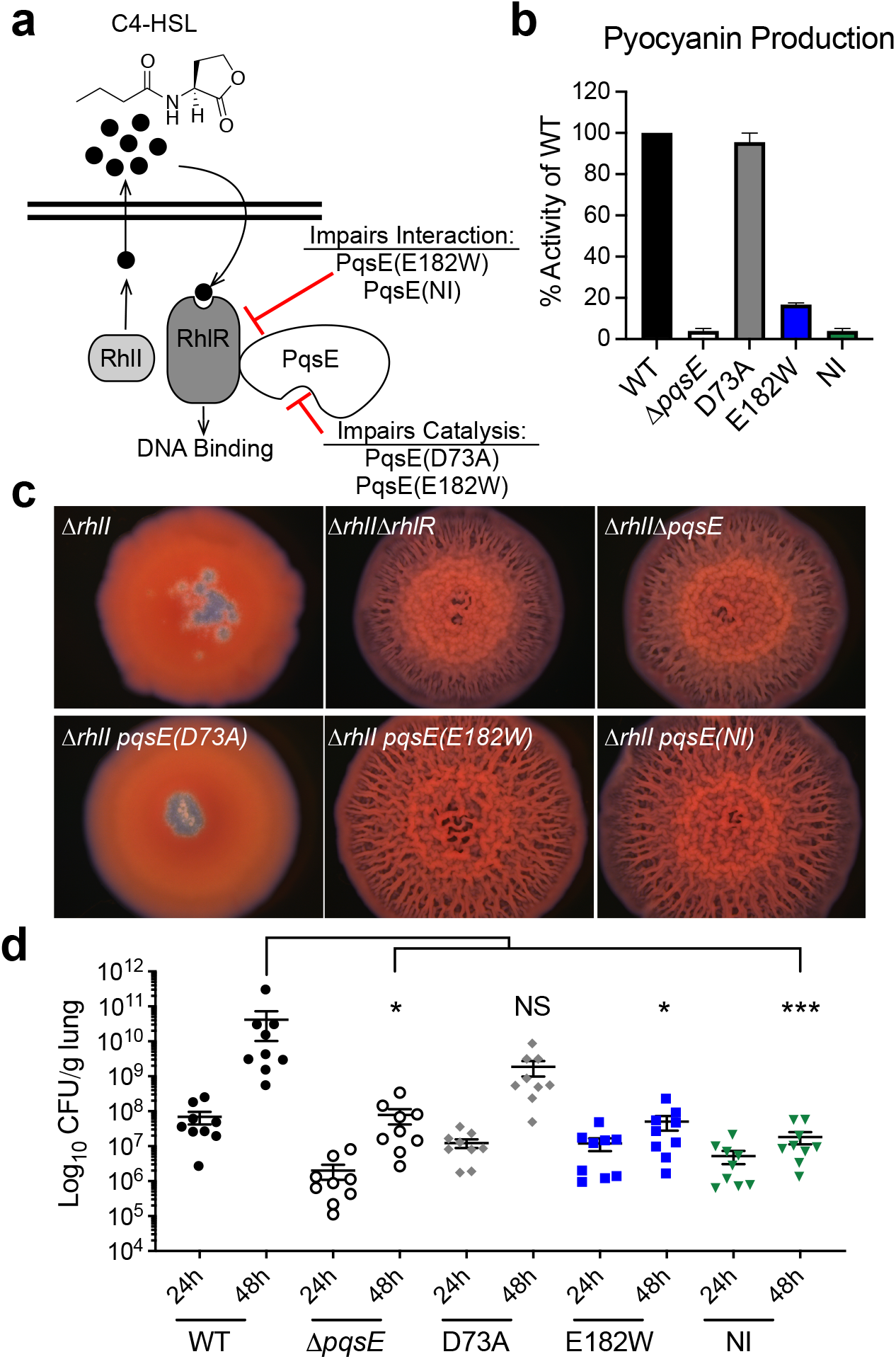
PqsE variants that are deficient in interaction with RhlR display attenuated pathogenicity. a) The Rhl QS system in *P. aeruginosa*. See text for details. b) Pyocyanin production by *P. aeruginosa* PA14 harboring the designated *pqsE* alleles at the native *pqsE* locus. The WT PA14 pyocyanin production level was set to 100%. Results are the average of two biological replicates. Error bars represent standard deviations. c) Colony biofilm morphologies of the designated *P. aeruginosa* PA14 strains. Biofilms were imaged after 5 d of growth at 7.78x magnification. Images are representative of three biological replicates. d) Lung colonization by the designated *P. aeruginosa* PA14, Δ*pqsE, pqsE(D73A), pqsE(E182W)*, or the *pqsE(NI)* strains in the murine infection model. Mice were euthanized at 24 and 48 h post-infection, and lung tissues were processed as described in Methods. All samples were plated for viable CFU on PIA. Each symbol represents the value for a single mouse. The data are pooled from two independent experiments. Error bars represent SEM. Statistical differences were determined using one-way ANOVA test and are as follows: ***, P < 0.001; *, P <0.05; NS means not significant.

We recently showed that the ability of PqsE to interact with RhlR can be weakened by introduction of an “inhibitor mimetic” mutation in which the glutamate residue at position 182 is altered to tryptophan (PqsE(E182W)) (20). The PqsE(E182W) variant is a stable protein that, while harboring all of the catalytic residues, nonetheless displays less than 10% of wildtype catalytic activity, as if an inhibitor is bound in the active site. Residue E182 is buried deep within the active site of PqsE, at a location distant from the three surface arginine residues (R243, R246, and R247) that are necessary for PqsE to interact with RhlR. Mutation of the PqsE R243, R246, and R247 residues to alanine completely abolishes the PqsE-RhlR interaction (21). When introduced into *P. aeruginosa*, neither PqsE(E182W) nor PqsE(R243A/R246A/R247A) (the latter called PqsE(NI) for “non-interacting”), can promote PqsE-RhlR-dependent virulence phenotypes, including production of the pyocyanin toxin. The effects of the PqsE E182W and PqsE NI substitutions on pyocyanin production are not through impairment of PqsE catalytic function, as the catalytically inactive variant, PqsE(D73A), remains fully capable of interacting with RhlR and driving pyocyanin production. These results indicate that PqsE has two independent functions, catalysis and interaction with RhlR, and it is interaction with RhlR, not catalysis, that is required for virulence.

From a drug discovery perspective, it is particularly promising that the PqsE active site E182W mutation weakens the distal PqsE-RhlR interaction, the consequence of which is suppression of virulence phenotypes. We assert this because a PqsE active site-targeting molecule would likely be able to bind with high affinity and be more amenable to medicinal chemistry than a molecule targeting the interaction site on the surface of the protein. Protein-protein interactions, which typically occur over large, shallow protein surfaces, have proven notoriously difficult to target with small molecules (22). Molecules that can bind in protein-protein interaction domains typically do so with weak affinity, are large and structurally complicated, and are difficult to optimize through medicinal chemistry efforts (23). With this notion in mind, in this study, we aimed to determine the mechanism by which the PqsE E182W mutation disrupts the PqsE-RhlR interaction, and whether it is possible to achieve a similar effect with a small molecule inhibitor that binds in the PqsE active site. In a proof of principle experiment, we use our PqsE variants to demonstrate that disrupting the PqsE-RhlR interaction indeed attenuates *in vivo P. aeruginosa* virulence in a mouse lung infection model. We employ crystallography to characterize the structure of the PqsE(E182W) protein. We probe the functions of the PqsE active site through mutagenesis. Finally, we present the next generation of PqsE active site-targeting small molecules for further synthetic optimization. Our results can inform the discovery and/or design of effective antimicrobial agents to treat *P. aeruginosa* infections.

## Results

### PqsE variants that cannot interact with RhlR display attenuated infection phenotypes in cell assays and in a mouse lung infection model

We have shown previously that, unlike overexpression of wildtype (WT) *pqsE*, overexpression of *pqsE* mutants encoding proteins that cannot interact with RhlR *in vitro* are impaired in promoting pyocyanin production in Δ*pqsE P. aeruginosa* PA14 (20,21). To verify that the PqsE variants of interest display similar defects in virulence phenotypes when the genes encoding them are expressed from the native locus, we constructed *P. aeruginosa* PA14 strains harboring *pqsE(D73A), pqsE(E182W),* and *pqsE(NI)* on the chromosome. Western blots showed that the WT and variant PqsE proteins were produced at the same levels and exhibited similar stabilities (Figure S1). WT *P. aeruginosa* PA14 made pyocyanin while the Δ*pqsE* strain did not (4% compared to WT). When the PqsE variant proteins were produced from the chromosomally-encoded genes, the results were entirely consistent with our previous findings for each PqsE variant produced from a plasmid. Specifically, the catalytically inactive PqsE(D73A) variant made nearly WT levels of pyocyanin (96%), the PqsE(E182W) inhibitor mimetic variant was severely impaired (17%), and the PqsE(NI) variant was incapable of driving pyocyanin production (4%) (Figure 1b).

Previously, we showed that the *P. aeruginosa* PA14 Δ*rhll* strain forms smooth colonies whereas the Δ*rhll ΔrhlR* and Δ*rhll* Δ*pqsE* double mutants form biofilms with hyper-rugose morphologies (24,25). Thus, both PqsE and RhlR are required to suppress hyper-rugose biofilm formation in the absence of the C4-HSL autoinducer. To determine which specific function of PqsE, catalysis and/or interaction with RhlR, is linked to the control of biofilm morphology, we tested our PqsE variants. Each *pqsE* mutant was incorporated at the native chromosomal locus in the Δ*rhl1* strain and biofilm morphology was assessed (Figure 1c). Both the Δ*rhlI pqsE(E182W)* and Δ*rhll pqsE(NI)* mutants formed hyper-rugose biofilms similar to those of the Δ*rhll* Δ*rhlR* and Δ*rhll* Δ*pqsE* mutants. Only the Δ*rhll* strain harboring the *pqsE(D73A)* mutation exhibited the smooth biofilm morphology of the parent Δ*rhll* strain. This result demonstrates that the PqsE-RhlR interaction controls biofilm morphology, and that PqsE catalytic activity is dispensable for this trait.

To explore the individual roles of PqsE catalysis and PqsE-RhlR interaction during host infection, we assessed the relative pathogenicity of WT *P. aeruginosa* PA14, Δ*pqsE, pqsE(D73A), pqsE(E182W)*, and the *pqsE(NI)* strains in a murine model of acute pneumonia. Mice were infected intratracheally with equal strain inoculum levels (~3×10^6^ CFU/mouse) and monitored over 48 h of infection. At 24 h, mice from all infection groups exhibited mild to moderate symptoms in response to infection, primarily displaying decreased mobility and increased breathing. Consistent with these symptoms, comparable levels of lung colonization were observed among all groups (Figure 1d). At 48 h, however, mice infected with either WT *P. aeruginosa* PA14 or the *pqsE(D73A)* mutant became more lethargic and appeared to progressively manifest additional symptoms, including increasingly labored breathing, hunched posture, and decreased response to stimuli. Moreover, those mice demonstrated >2 log increase in the bacterial burden compared to 24 h. In stark contrast, mice infected with the Δ*pqsE, pqsE(E182W)*, or *pqsE(NI)* strains continued to display mild clinical symptoms with almost no change in their lung bacterial burden (Figure 1d). Together, the above findings demonstrate that the PqsE-RhlR interaction, and not PqsE catalytic activity, is responsible for shaping the pathogenicity of *P. aeruginosa* PA14 *in vitro* and *in vivo*.

### The PqsE E182W substitution induces a loop rearrangement near the active site

The above murine infection experiment demonstrated that weakening the ability of PqsE to interact with RhlR by mutating an active site residue (PqsE(E182W)) causes a similarly severe reduction in infectivity to that caused by complete elimination of the PqsE-RhlR interaction (PqsE(NI)). To understand, at an atomic level, what conformational changes the E182W alteration induced in PqsE to affect its ability to interact with RhlR, we determined the structure of PqsE(E182W) (Figure 2a). Although the PqsE(E182W) crystals grow in the same P_3_221 crystal form as WT PqsE, there is significant structural rearrangement in the active site. In WT PqsE, the sidechain of E182 lies at the edge of the ligand binding site and makes hydrogen bonds with R191 and Q272. These interactions do not occur in PqsE(E182W), and rather, new interactions are made between the sidechain of the introduced W182 residue and F276, L277, and P278. The E182W change induces the rearrangement of the G270-L281 loop between helices 6 and 7, with a portion of that loop becoming disordered. Examination of electron density within the active site indicated a surprising consequence - with the sidechain of E280 relocating by 12 Å and becoming Fe bound in the center of the active site. Indeed, the repositioned E280 sidechain directly binds both Fe atoms, acting as a bridging ligand, and thus is most likely responsible for the stabilizing effect the E182W alteration has on PqsE, which we have reported previously (20).

**Figure 2:**
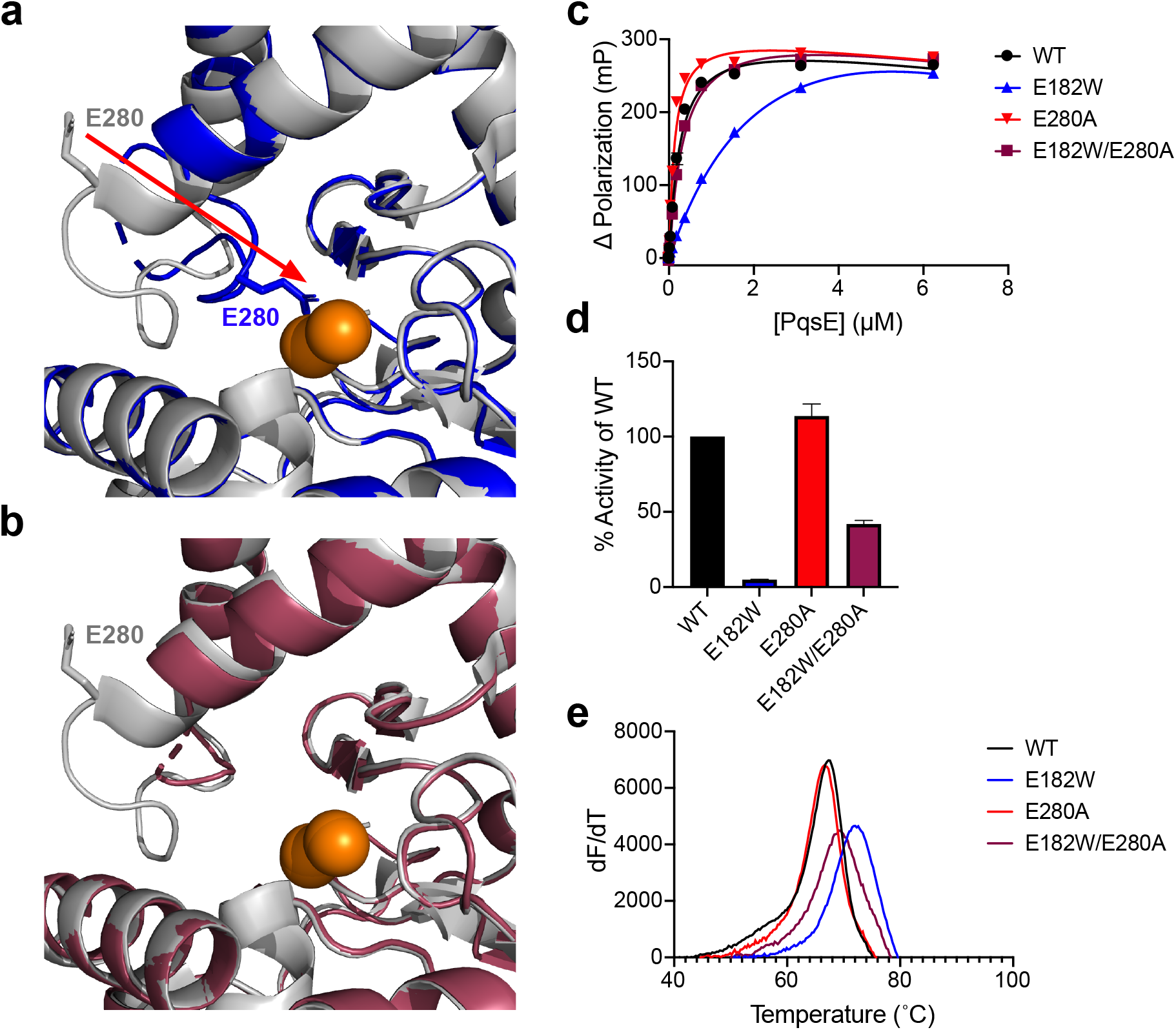
The PqsE E182W substitution induces a loop rearrangement that repositions residue E280. a) Structure of PqsE(E182W) (blue) overlaid with that of WT PqsE (PDB: 2Q0I, gray). The active site iron atoms are in shown in orange. The red arrow indicates repositioning of E280 in PqsE(E182W) relative to its position in WT PqsE. b) Structure of PqsE(E182W/E280A) (maroon) overlaid with that of WT PqsE (as in a). In panels a and b, residue E280 is shown in stick representation. c) Binding of WT and variant PqsE proteins to the active site fluorescent probe BB562. K_app_ was determined in two independent experiments performed in triplicate. d) Hydrolysis of 4-methylumbelliferyl butyrate by the designated purified PqsE proteins. Values are represented as % activity of WT PqsE protein. Results are the average of two independent experiments performed in triplicate. Error bars represent standard deviations. e) First derivative plots (dF/dT is defined as the change in SYPRO Orange fluorescence divided by change in temperature) of melting curves for the designated PqsE proteins. The peak of each curve is defined as the T_m_ of that protein.

To test whether repositioning of PqsE residue E280 underpinned the inhibitor mimetic characteristics of the PqsE(E182W) protein, we engineered the E280A substitution into PqsE(E182W) to make PqsE(E182W/E280A). We determined the crystal structure of PqsE(E182W/E280A) revealing further changes in the PqsE active site. In this case, the presence of an alanine at residue 280 eliminated interaction with the iron atoms, and the loop consisting of residues D275-L281 was disordered (Figure 2b). The remainder of the loop was ordered and its structure reverted to a conformation that was more native-like than in PqsE(E182W). Consequently, the removal of the glutamate-Fe interaction via the E280A substitution apparently unblocked the active site of PqsE, partially restoring WT activity. Indeed, whereas PqsE(E182W) exhibits reduced binding affinity for the active site fluorescent probe, BB562, PqsE(E182W/E280A) displays WT binding affinity for the probe (Figure 2c). Furthermore, PqsE(E182W) has reduced hydrolytic capacity for a synthetic ester substrate, with only 7% activity relative to WT PqsE. The catalytic function of the PqsE(E182W/E280A) variant was greatly improved by the “unblocking” of the active site (34% compared to WT PqsE, Figure 2d). Consistent with the repositioning of E280 into the PqsE active site contributing to the increased stability of the PqsE(E182W) protein relative to WT PqsE, introduction of the E280A substitution reduced the T_m_ of the PqsE(E182W/E280A) protein to nearly that of WT PqsE (Figure 2e).

Introduction of the E280A alteration into PqsE(E182W) partially corrects the defects in small molecule binding and hydrolysis. Nonetheless, the PqsE(E182W/E280A) variant remains incapable of enhancing RhlR transcription factor activity in an *Escherichia coli* reporter assay and it does not drive pyocyanin production in *P. aeruginosa* PA14 (Figure 3a and 3b, respectively). Likewise, PqsE(E182W/E280A) is not improved for interaction with RhlR *in vitro* (Figure 4a). These findings further demonstrate the independence of the PqsE catalytic and virulence functions. Curiously, neither the PqsE(E182W) nor the PqsE(E182W/E280A) structure showed any changes in the region of the protein that interacts with RhlR (R243, R246, and R247 on helix 5, Figure 2a and 2b, respectively). This result preliminarily suggests that either PqsE possesses an additional region of interaction with RhlR, perhaps employing the G270-L281 loop that is rearranged in PqsE(E182W) or partially disordered in PqsE(E182W/E280A), or alternatively, that a particular conformation of the G270-L281 loop is required to allosterically promote the interaction with RhlR. Irrespective of the underlying mechanism, the above structural and biochemical analyses characterizing PqsE(E182W/E280A) show that it is possible to disrupt the PqsE-RhlR interaction by manipulating the integrity of the G270-L281 loop that forms one face of the PqsE active site.

**Figure 3:**
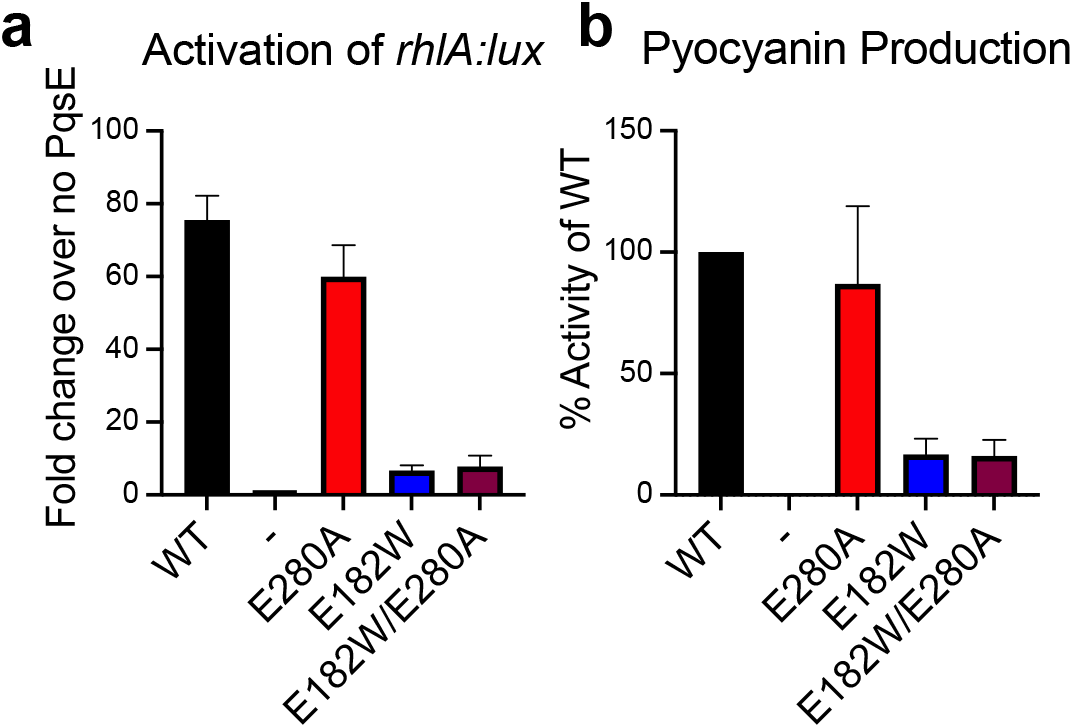
The PqsE E280A substitution does not restore activation of RhlR transcriptional activity or pyocyanin production. a) Light production from *E. coli* carrying *P_BAD_-rhlR, PrhlA-luxCDABE*, and the designated *pqsE* allele on the pACYC184 plasmid was measured following growth in the presence of 100 nM C4-HSL. The “-” symbol represents the strain carrying the empty pACYC184 vector. Results are the average of two biological replicates performed in technical triplicate. b) Pyocyanin production was measured from Δ*pqsE P. aeruginosa* PA14 carrying the designated *pqsE* allele on the pUCP18 plasmid. The “-” symbol represents the strain carrying an empty pUCP18 plasmid. Results shown are the average of two biological replicates. Pyocyanin production from the strain with WT PqsE was set to 100%. Error bars represent standard deviations.

**Figure 4:**
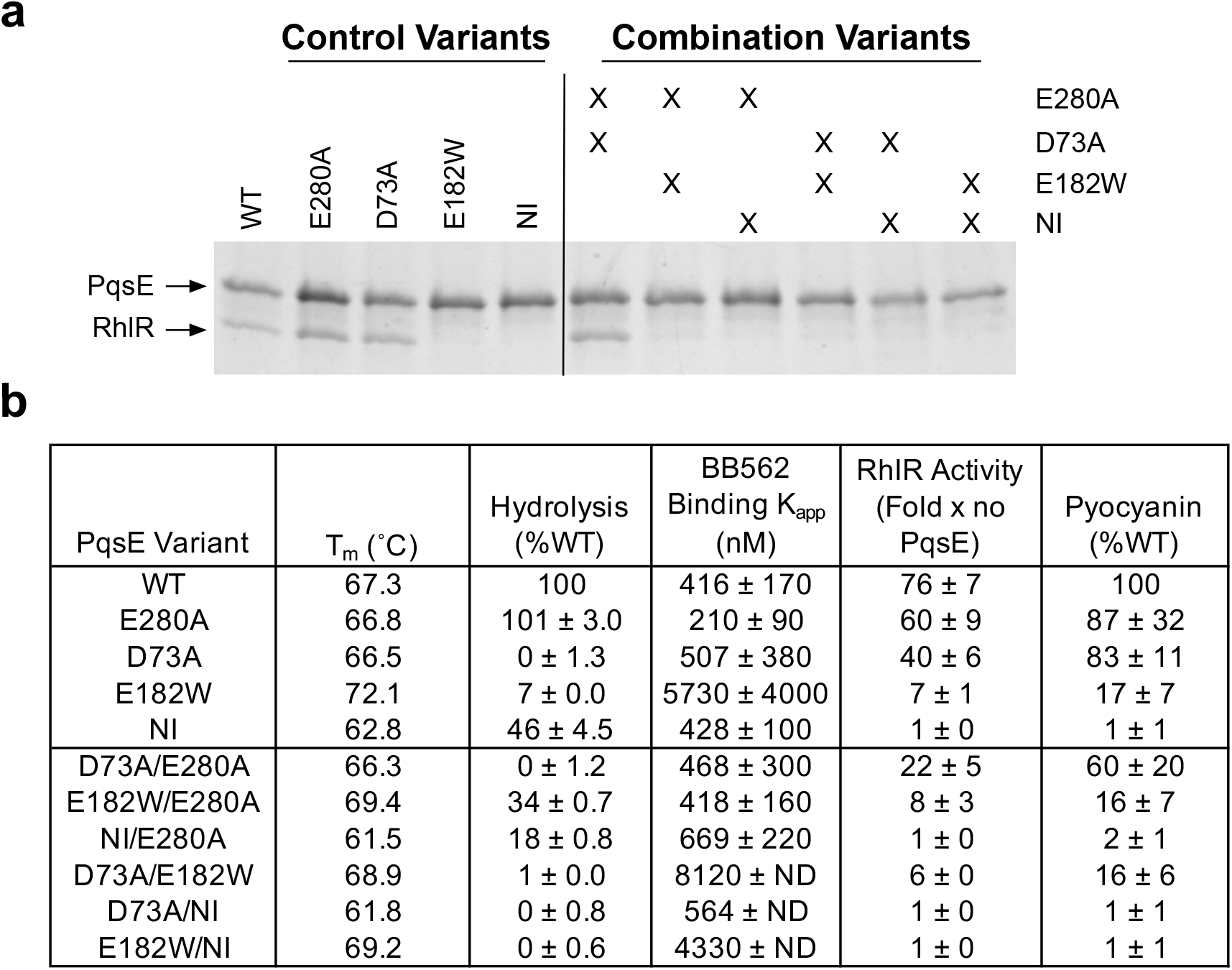
PqsE enzymatic activity is dispensable for RhlR interaction, RhlR transcriptional activity, and pyocyanin production. a) SDS-PAGE analysis of protein complexes formed in *in vitro* pull-down assays. 6xHis-PqsE proteins were immobilized on Ni resin and exposed to lysate containing RhlR. The resulting protein complexes were washed and eluted from the Ni resin and loaded into the lanes of the gel. PqsE appears as a ~34 kDa band and RhlR as a ~28 kDa band. For the variant combinations, “X” denotes that the protein in that lane contains the designated substitutions listed to the right. b) Measured activities of PqsE variants in the designated assays. The full data from these analyses are provided in Figures S3 and S4 including replicate and statistical information. “ND” denotes that error could not be calculated for the binding curve, which occurred when the binding curve was too shallow or did not achieve saturation.

### PqsE variants harboring combinations of substitutions decouple catalytic activity from RhlR interaction

Our previous and above work comparing the inhibitor mimetic and RhlR non-interacting PqsE variants to the catalytically inactive PqsE(D73A) protein strongly suggest that the ability of PqsE to interact with RhlR and, in turn, drive virulence phenotypes is independent of PqsE enzyme activity. To confirm this notion, and, additionally, to explore the potential for influence of one activity on the other, we constructed PqsE variants containing combinations of amino acid substitutions underlying defects in enzyme activity, RhlR interaction, or both functions. Purity of these PqsE variant proteins was verified by SDS-PAGE analysis (Figure S2). We assessed the phenotypes of this set of variants *in vitro*, in recombinant *E. coli*, and in *P. aeruginosa* PA14. Specifically, we measured interaction of the purified PqsE variant proteins with RhlR by our pull-down assay (Figure 4a), intrinsic stability by T_m_ measurements (differential scanning fluorimetry, DSF), catalytic function by hydrolysis of the synthetic ester substrate 4-methylumbelliferyl butyrate (MU-butyrate), and active site accessibility via binding of the fluorescent active site probe, BB562 (Figure 4b). We measured enhancement of RhlR transcriptional activity in the *E. coli* reporter assay and virulence by pyocyanin production in *P. aeruginosa* PA14 (Figure 4b). WT PqsE and the variants PqsE(E280A), PqsE(D73A), PqsE(E182W), and PqsE(NI) have all been previously characterized in this suite of assays and were included in our analyses for comparison to the variants containing combinations of substitutions. As shown in Figure 4 (and Figures S3 and S4), the ability of PqsE to form a complex with RhlR always tracked with activation of RhlR transcription factor activity and pyocyanin production. By contrast, enzymatic capability did not correlate with the ability to form a complex with RhlR, activate RhlR transcription, or to produce pyocyanin.

One potential complication in the above analyses is that PqsE proteins harboring the “NI” triple arginine substitutions are less stable than WT and other of our variant PqsE proteins when overproduced in cells from a plasmid. Therefore, all variant proteins containing the NI substitutions were detected at lower levels compared to the other PqsE variant proteins in both *E. coli* and *P. aeruginosa* PA14 cell lysates (Figures S5 and S6). This feature could potentially have been the source of their apparent reduced activities in the RhlR transcriptional reporter assay and the pyocyanin assay. Our results with the PqsE(E182W/NI) variant show, however, that this is not the case. Due to the stabilizing effect of the E182W alteration (Figures 2 and S3), the PqsE(E182W/NI) protein was produced and detected at WT levels in both *E. coli* and *P. aeruginosa* lysates (Figures S5 and S6). Nonetheless, PqsE(E182W/NI) was completely inactive in both the pyocyanin production and RhlR transcriptional activity cell-based assays (Figure 3). This result confirms that the inability of select PqsE variants to activate RhlR and drive pyocyanin production stems from loss of the PqsE-RhlR interaction, and is not the result of decreased PqsE protein production or stability.

### Synthetic optimization of an active site-targeting small molecule scaffold

The results of our structural and mutagenic analyses suggest that manipulation of the PqsE active site, such that the G270-L281 loop becomes rearranged or disordered will result in decreased *P. aeruginosa* virulence due to inhibition of the PqsE-RhlR interaction. Thus, it is of interest to develop molecules that bind in the PqsE active site and, in so doing, inhibit the PqsE-RhlR interaction. We previously characterized two active site-targeting PqsE inhibitors, BB391 and BB393 (20). Crystallographic analyses of each compound bound to PqsE revealed their respective binding poses and interactions in the active site. Each inhibitor exhibited mid-nanomolar competition with the BB562 active site probe for binding to PqsE (20).

Guided by our crystal structures of BB391 and BB393 bound to PqsE, we designed a series of BB391-BB393 hybrid derivatives to probe the contributions of each moiety to binding affinity (Figure 5a). BB580 and BB581 were designed as the core hybrid structures, in which BB580 maintained the same amide bond orientation as in BB391, and BB581 possessed the amide bond orientation of BB393 (see position “1” in Figure 5a). BB580 outcompeted the BB562 probe with an EC_50_ of 99 nM compared to BB581 which exhibited an EC_50_ of 281 nM (Figure 5b,c). This result demonstrates that preservation of the BB391 amide bond orientation coupled with the indazole ring, also from BB391, is superior for proper positioning of the carbonyl oxygen to form a hydrogen bond with PqsE residue S285. The structure of BB391 bound to PqsE showed that the possibility existed for pi-stacking between the core phenyl ring and the H71 residue. Derivatives BB585, BB586, and BB589 were designed to test enhancement of pi stacking with alternative aromatic groups at this core position (see position “2” in Figure 5a). None of these three derivatives, featuring thiazole, pyrazole, or pyridine rings, respectively, bound as tightly as BB580, and therefore among the molecules tested, a phenyl ring was deemed ideal at this core position (Figure 5b,c). Finally, derivatives BB582, BB583, BB584, BB587, and BB588 were designed to assess the importance of a hydrophobic moiety on the BB393 molecule (see position “3” in Figure 5a), which in the crystal structure, nestles into a hydrophobic groove near the solvent-exposed entrance to the PqsE active site. The results with BB587 and BB588 show that elimination of either the methyl or ethyl groups, respectively, leads to slightly decreased binding affinity (EC_50_ = 273 nM and 290 nM, respectively). However, if the morpholine ring is removed and either a phenyl (BB583) or tert-butyl (BB584) group is installed, a modest increase in binding affinity is achieved (EC_50_ =71 nM and 34 nM, respectively) (Figure 5b,c). Such improvement was not observed for BB582, featuring a pyrazine ring, suggesting that increased hydrophobicity of this portion of the molecule tracks with increased engagement of the hydrophobic groove in PqsE. None of the derivatives described here inhibit the PqsE-RhlR interaction. Within that context, BB584 displayed the tightest binding to PqsE, and provides a starting scaffold for the design of new derivatives.

**Figure 5:**
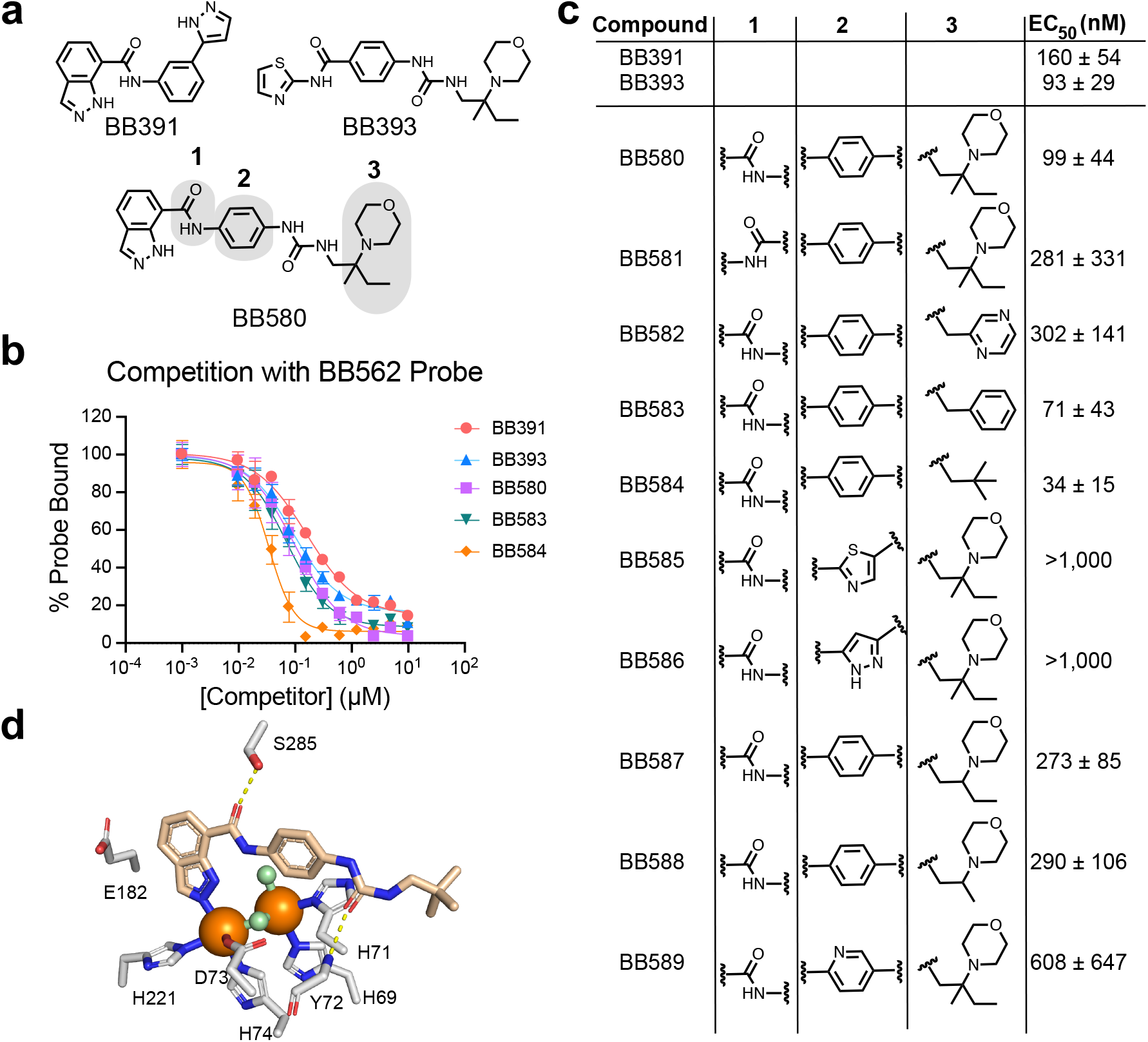
Optimization of new PqsE active site-targeting small molecules. a) Structures of the precursor molecules BB391 and BB393 and the hybrid derivative BB580. Positions denoted 1, 2, and 3 were derivatized to explore binding affinity for the PqsE active site. b) Fluorescence polarization competition curves for select BB391-BB393 hybrid derivatives competing with the BB562 active site probe for binding to PqsE. Polarization value for PqsE-BB562 in the absence of competitor is defined as 100% Probe Bound. All polarization values were background-subtracted by the reading for the probe in the absence of PqsE (background fluorescence). c) Competitive fluorescence polarization EC_50_ values calculated for all BB391-BB393 derivatives, determined from one experiment performed in triplicate. d) Structure of BB584 bound to PqsE. Both the BB584 molecule and key PqsE amino acid sidechains are shown as sticks. Amino acid sidechain carbons are depicted in gray and BB584 carbons are shown in tan. Iron atoms and water molecules are shown as orange and green spheres, respectively. Oxygen and nitrogen atoms are in red and blue, respectively. Hydrogen bonds are shown as dotted yellow lines.

We determined the crystal structure of BB584 bound to PqsE, which showed a similar orientation in the active site and similar ligand-protein interactions to those observed in the crystal structures of PqsE with BB391 and BB393 bound in the active site (Figure 5d). Specifically, the nitrogen on the indazole ring of BB584 bonds with the Fe2 atom, displacing a water molecule that normally resides at this position. Two additional hydrogen bonds exist between BB584 and PqsE sidechains: the BB584 amide oxygen with the hydroxyl group of the S285 sidechain, and the BB584 urea oxygen with the backbone amide N-H of residue Y72. As predicted, the tert-butyl group tucks into the hydrophobic groove that sits between the *α*-helix consisting of PqsE residues S104-L116 and the backbone of the conserved ^69^HXHXDH^74^ motif. Analogous to the structures of BB391 and BB393 bound to PqsE, no significant conformational changes in the protein were induced by binding of BB584 (RMSD of 0.24 Å for 293 Ca atoms), possibly explaining why none of the three compounds disrupt the PqsE-RhlR interaction. Nonetheless, the structures of these ligands bound to WT PqsE along with the structures of PqsE(E182W) and PqsE(E182W/E280A) can be used as guides in the design of new derivatives with the potential to disrupt the interaction between PqsE and RhlR.

## Discussion

PqsE has two activities, a protein-protein interaction with RhlR, which increases RhlR transcriptional activity at target promoters, and an esterase activity, the substrate and product of which are currently unknown. We showed that these two functions are separable. Moreover, we engineered a set of PqsE variant proteins that possess every combination of the two activities (catalytic^+^/interaction^+^, catalytic^-^/interaction^+^, catalytic^+^/interaction^-^, and catalytic^-^/interaction^-^). Analysis of these proteins, coupled with previous work, shows that the PqsE-RhlR interaction is linked to virulence factor production in *P. aeruginosa* PA14. Here, we demonstrated that the PqsE-RhlR interaction also controls development of *P. aeruginosa* PA14 biofilm morphology and is crucial for establishing an infection in a host animal. Remarkably, PqsE catalytic function is dispensable for regulation of RhlR-transcriptional activity, biofilm morphology, virulence factor production, and animal infectivity. As the PqsE variant phenotypes are consistent between all assays performed, the set of cell-based assays we employ here can effectively be used in lieu of infectivity assays in animals to probe and predict the potential of small molecules as *P. aeruginosa* antibiotics that target PqsE.

Our finding that the PqsE-RhlR interaction was disrupted following introduction of the E182W substitution in the PqsE active site was surprising given that this residue is distal to the RhlR interaction site (21). The crystal structure of PqsE(E182W) revealed that a loop rearrangement inserts the E280 residue into the active site to coordinate both of the iron atoms at this site. It is the repositioning of E280 in the PqsE(E182W) variant – blocking substrate access to the catalytic iron atoms – that is responsible for the “inhibitor mimic” nature of this variant. Catalytic activity is restored in the PqsE(E182W/E280A) variant because the alanine substitution clears the active site enabling substrate binding, however, PqsE(E182W/E280A) remains incapable of interacting with RhlR (Figures 3 and 4). Analysis of the PqsE(E182W/E280A) structure, and the surprisingly modest conformational changes that are apparent compared to the structures of WT PqsE and PqsE(E182W) (Figure 2a,b), suggest that the structural integrity of the G270-L281 loop is essential for PqsE to interact with RhlR. These results confirm that, although the RhlR-interaction is independent of PqsE catalytic activity, structural changes induced through the active site of PqsE can disrupt interaction with RhlR.

All prior crystal structures of WT PqsE showed that the E280 residue resides on the surface with the glutamate sidechain directed outward into the solvent. Our data show that single substitution of this glutamate with alanine (i.e., PqsE(E280A)) increased accessibility of the PqsE active site (Figure 2c). This finding potentially points to naturally occurring PqsE conformational dynamics in which the position of the E280 sidechain alternates between facing the solvent and being inserted into the active site. While not proven, it is possible that E280 serves as a dynamic gate-keeper residue for the active site. Perhaps, its position determines whether PqsE will undergo catalysis and/or will interact with RhlR. Such a mechanism would make considerations of the PqsE E280 sidechain position key for future drug discovery efforts (26).

Figure 5 presents a new series of molecules, inspired by the previous PqsE inhibitors BB391 and BB393, all of which bind in the PqsE active site. Our competitive binding assay allowed us to rank derivatives by their relative affinities providing preliminary structure-activity relationships. Among this set, compound BB584 binds most tightly in the PqsE active site (EC_50_ = 34 nM). The crystal structure of the PqsE-BB584 complex solved here (Figure 5c) provides needed information for launching structure-guided design of the next generation of molecules targeting the PqsE active site. None of the inhibitors characterized here perturb the PqsE-RhlR interaction. However, our PqsE variant analysis and companion structures make it clear that it should be possible to design high affinity active site-targeting compounds that disrupt the PqsE-RhlR interaction via an allosteric mechanism, presumably involving movement of the G270-L281 loop.

## Methods

### Strains, Media, and Molecular Procedures

The *P. aeruginosa* UCBPP-PA14 strain was used as the parental strain for all experiments involving *P. aeruginosa*. All strains were grown in Luria-Bertani broth, unless otherwise stated, and antibiotics were used at the following concentrations: ampicillin (200 μg/mL), kanamycin (100 μg/mL), tetracycline (10 μg/mL), carbenicillin (400 μg/mL), gentamycin (30 μg/mL), and irgasan (100 μg/mL). Plasmids were constructed following a previously reported site-directed mutagenesis protocol (27) and were transformed into *P. aeruginosa* PA14 strains as described (28). Deletion of and point mutations in *pqsE* were generated by a previously reported method, with some modifications (24). Briefly, *pqsE* variants were cloned onto the pEXG2 vector. *E. coli* SM10λ*pir* carrying each pEXG2-*pqsE*-containing plasmid was mated with *P. aeruginosa* PA14 or the Δ*rhll* strain and exconjugants were selected on LB agar containing gentamycin and irgasan. Colonies were grown in LB medium at 37 °C for 1-2 h and plated on LB agar containing 5% sucrose to force elimination of the *sacB* gene on the plasmid. Resulting colonies were patched onto LB agar plates and onto plates containing gentamycin, and *pqsE* variants in Gent^S^ colonies were confirmed by sequencing. Strains used in this study are listed in Supplemental Table S1.

### General Methods

6xHis-PqsE proteins were purified for biochemical assays (tagged) and crystallography (tag removed) as described previously (20). Enzyme activities of purified 6xHis-PqsE variants were measured as previously reported with 4-methylumbelliferyl butyrate (MU-butyrate) as the substrate (21). Fluorescence polarization assays with the BB562 active site probe were performed as described (20). The melting temperature (T_m_) of each purified 6xHis-PqsE variant was measured using differential scanning fluorimetry (DSF) as described previously (20). Interaction between PqsE and RhlR proteins *in vitro* was measured as described previously using pull-down assays (20). Pyocyanin production by *P. aeruginosa* PA14 strains carrying *pqsE* variants on the pUCP18 plasmid was measured as described (20). The ability of PqsE variants to increase RhlR transcription factor activity in an *E. coli* reporter assay was assessed as previously described (20). All small molecule syntheses and characterization are described in the Supplementary Information.

### Colony Biofilm Morphology Assay

Cultures were grown overnight in LB broth with shaking at 37 °C and 1 μL of culture was spotted onto a 60 x 15 mm Petri plate containing 10 mL biofilm medium (1% Tryptone, 1% agar, 40 mg/L Congo Red, 20 mg/L Coomassie Brilliant Blue). Biofilms were grown at 25 °C for several days and imaged throughout their development on a Leica stereomicroscope M125 mounted with a Leica MC170 HD camera at 7.78x magnification.

### Mouse Lung Infection Studies

For all mouse experiments, *P. aeruginosa* strains were grown on *Pseudomonas* Isolation Agar (PIA) for 16–18 h at 37°C and suspended in PBS to an OD_600_ of 0.5, corresponding to ~10^9^ CFU/mL. These samples were adjusted spectrophotometrically and then diluted to the appropriate OD_600_ in phosphate-buffered saline (PBS). Eight-to-ten-week-old female BALB/c mice (Jackson Laboratories, Bar Harbor, ME) were anesthetized by intraperitoneal (i.p.) injection of 0.2 mL of a mixture of ketamine (25 mg/mL) and xylazine (12 mg/mL). Mice were infected by noninvasive intratracheal instillation (29) of 50 μL of ~3 × 10^6^ CFU of *P. aeruginosa* WT or isogenic mutants. Mice were euthanized at 24 and 48 h post-infection and whole lungs were collected aseptically, weighed, and homogenized in 1 mL of PBS. Bacterial loads in tissue homogenates were enumerated by serial dilution and plating on PIA. Comparison and analyses of the numbers of viable bacteria obtained in lung homogenates were performed using GraphPad Prism version 7 software. Results were analyzed using one-way analysis of variance (ANOVA) and were compared using the Kruskal-Wallis test for comparison of three groups or the Mann-Whitney *U* test for analysis of two groups.

### Protein Crystallography

Purified proteins (~10 mg/mL) and protein-compound complexes were crystallized by hanging drop vapor diffusion at 22 °C following mixing at a 1:1 ratio with well buffer. Crystallization buffers used were as follows: PqsE(E182W) (0.1 M HEPES pH 7.5, 0.2 M MgCl_2_, 15% (w/v) PEG 400) cryoprotected with 20% (v/v) glycerol prior to freezing, PqsE(E182W/E280A) (0.1 M HEPES pH 7.5, 0.2 M MgCl_2_, 27% (w/v) PEG 400) cryoprotected with 10% (v/v) ethylene glycol prior to freezing, and PqsE-BB584 (0.1 M HEPES pH 7.5, 0.2 M MgCl_2_, 15% (v/v) 2-propanol) cryoprotected with 30% (v/v) ethylene glycol with additional BB584 (50 μM) in the cryoprotectant solution. Crystals typically formed within 48 h, but in some cases, took up to 5 days to form. PqsE(E182W), PqsE(E182W/E280A), and PqsE-BB584 crystals each grew in the same trigonal space group as had been previously observed for WT PqsE, PqsE-BB391 and PqsE-BB393 (P_3_21, a=b= 60 Å c= 146 Å α=β=90°, γ=120°) with one molecule in the asymmetric unit. Data were collected on either the 17-ID-1 beamline of the NSLS-II synchrotron (PqsE(E182W) and PqsE(E182W/E280A)), or on a Rigaku MicroMax 007HF rotating anode source (PqsE-BB584) (Table S2). Data were processed either with XDS (30) and AIMLESS (31) or with DENZO and SCALEPACK (32). The starting model for each refinement was a prior ligand-bound structure (7KGX, (20)) with the ligand and water molecules removed but the iron atoms retained, subjected to rigid-body refinement followed by conventional refinement using Phenix.refine (33). The structures were iteratively rebuilt using Coot (34) and refined with Phenix.refine. Both PqsE variant proteins exhibited significant conformational changes in a loop region (G270-L281) relative to the starting model, which remodeled the active site. In the case of the BB584 ligand, an atomic model of the ligand was fit to the difference electron density observed in the active site of PqsE and refined with partial occupancy (0.93). Final refinement statistics are shown in Table S2 including the identifiers for the structures’ depositions in the Protein Data Bank (PqsE(E182W) PDB ID: 7TZ9, PqsE(E182W/E280A): 7U6G, and PqsE-BB584 PDB ID: 7TZA).

## Supporting information

All supplemental figures and text

## Ethics Statement

All mouse procedures were performed in accordance with the established guidelines of the Emory University Institutional Animal Care and Use Committee (IACUC) under protocol number DAR-201700441. This study was carried out in strict accordance with established guidelines and policies at Emory University School of Medicine, the recommendations in the Guide for Care and Use of Laboratory Animals (35), as well as local, state, and federal laws.

## Acknowledgements

We thank members of the Bassler laboratory for helpful advice and discussions. We are grateful to William Miller and Brad Henke for guidance in chemical strategies and synthetic procedures. Molecules used in this study were synthesized at WuXi AppTec. Protein crystallography was performed in the Macromolecular Crystallography Core Facility at Princeton University. The AMX beamline of the National Synchrotron Light Source II, a U.S. Department of Energy (DOE) Office of Science User Facility operated for the DOE Office of Science by Brookhaven National Laboratory was also used under Contract DE-SC0012704. This work was supported by the Howard Hughes Medical Institute, NIH grant 2R37GM065859, and National Science Foundation grant MCB-2043238 to B.L.B., and NIH grant F32GM134583 to I.R.T. The content herein is solely the responsibility of the authors and does not represent the official views of the National Institutes of Health. The authors declare that they have no competing financial interests.

## Author Contributions

I. R.T, P.D.J., and D.A.M. conducted experiments; I.R.T, P.D.J., D.A.M., J.B.G., and B.L.B. designed experiments and prepared the manuscript.

