## Supplementary material for "The PqsE active site as a target for small molecule antimicrobial agents against *Pseudomonas aeruginosa*": All supplemental figures and text

**This PDF file includes:**

Figures S1 to S7
Tables S1 and S2
Supplementary text

### Abstract

The opportunistic pathogen *Pseudomonas aeruginosa* causes antibiotic resistant, nosocomial infections in immuno-compromised individuals, and is a high priority for antimicrobial development. Key to pathogenicity in *P. aeruginosa* are biofilm formation and virulence factor production. Both traits are controlled by the cell-to-cell communication process called quorum sensing (QS). QS involves the synthesis, release, and population-wide detection of signal molecules called autoinducers. We previously reported that activity of the RhlR QS transcription factor depends on a protein-protein interaction with the hydrolase, PqsE, and PqsE catalytic activity is dispensable for this interaction. Nonetheless, the PqsE-RhlR interaction could be disrupted by substitution of an active site glutamate residue with tryptophan (PqsE(E182W)). Here, we show that disruption of the PqsE-RhlR interaction via either the E182W change or alteration of PqsE surface residues that are essential for the interaction with RhlR, attenuates *P. aeruginosa* infection in a murine host. We use crystallography to characterize the conformational changes induced by the PqsE(E182W) substitution to define the mechanism underlying disruption of the PqsE-RhlR interaction. A loop rearrangement that repositions the E280 residue in PqsE(E182W) is responsible for the loss of interaction. We verify the implications garnered from the PqsE(E182W) structure using mutagenic, biochemical, and additional structural analyses. We present the next generation of molecules targeting the PqsE active site, including a structure of the tightest binding of these compounds, BB584, in complex with PqsE. The findings presented here provide insight for drug discovery against *P. aeruginosa* with PqsE as the target.

### Summary

The human pathogen *Pseudomonas aeruginosa* is resistant to many currently used antibiotics, making it a burden of urgent clinical importance. *P. aeruginosa* pathogenicity is controlled by the bacterial cell-to-cell communication process called quorum sensing (QS). The function of one protein that controls *P. aeruginosa* QS-directed virulence, RhlR, requires a protein-protein interaction with an enzyme called PqsE. When PqsE is blocked from interacting with RhlR, *P. aeruginosa* is avirulent and incapable of infecting an animal host. Here, we validate the PqsE-RhlR interaction as a target for antibiotic development, and we present a mechanism for how such antibiotics could disrupt the PqsE-RhlR interaction. Discovery of new antibiotics would fulfill an unmet healthcare need by providing treatments to combat *P. aeruginosa* infections.

**Supplementary Figures**

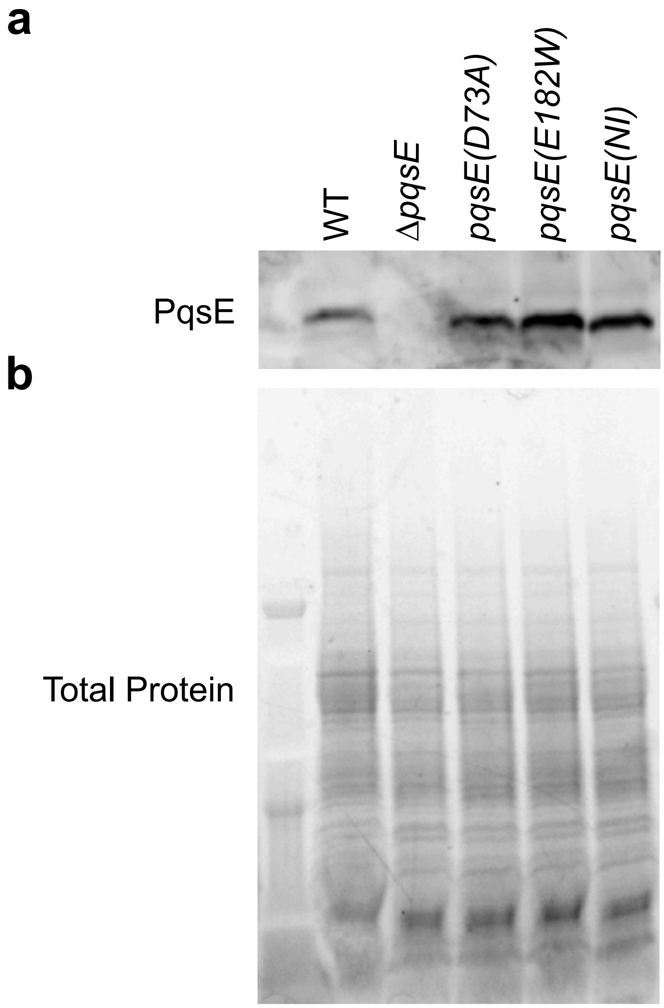

**Fig. S1.** Western blot analysis of PqsE proteins made by *P. aeruginosa* PA14. *P. aeruginosa* PA14 harboring the designated *pqsE* alleles at the native *pqsE* locus were grown to $OD_{600}$ = 1.5, lysed, and subjected to SDS-PAGE and Western blot analysis. a) Western blot of the gel in panel b following incubation with a primary rabbit polyclonal $\alpha$-PqsE antibody and secondary $\alpha$-rabbit HRP-conjugated antibody. Detection was via chemiluminescence. b) Total protein was imaged using a UV-activated in-gel protein stain.

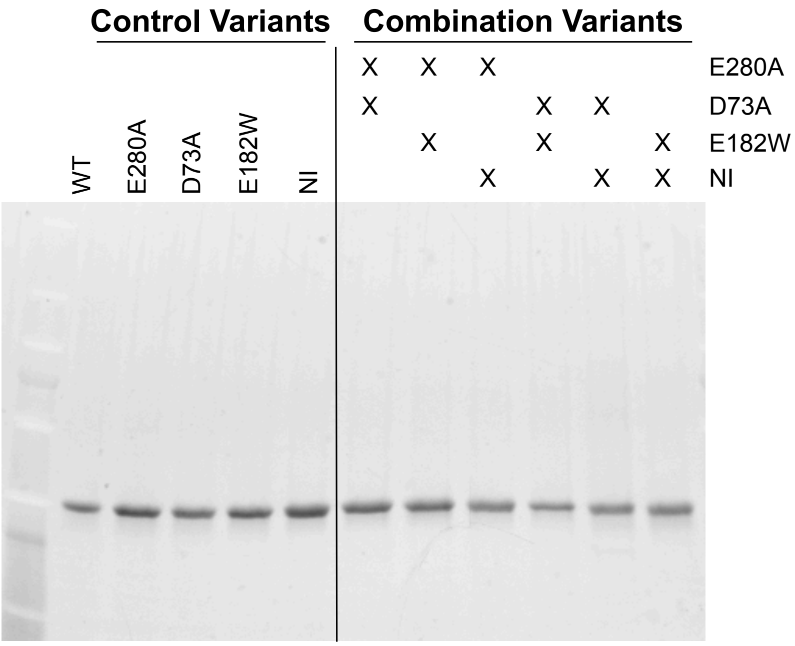

**Fig. S2.** SDS-PAGE analysis of purified PqsE proteins. Approximately 0.6 µg of the specified purified PqsE protein was loaded into each lane and imaged using a UV-activated in-gel protein stain. For the combination variants, “X” signifies that the protein in that lane harbors the corresponding substitution indicated on the right. All purified PqsE proteins appeared as ~34 kDa bands on the gel.

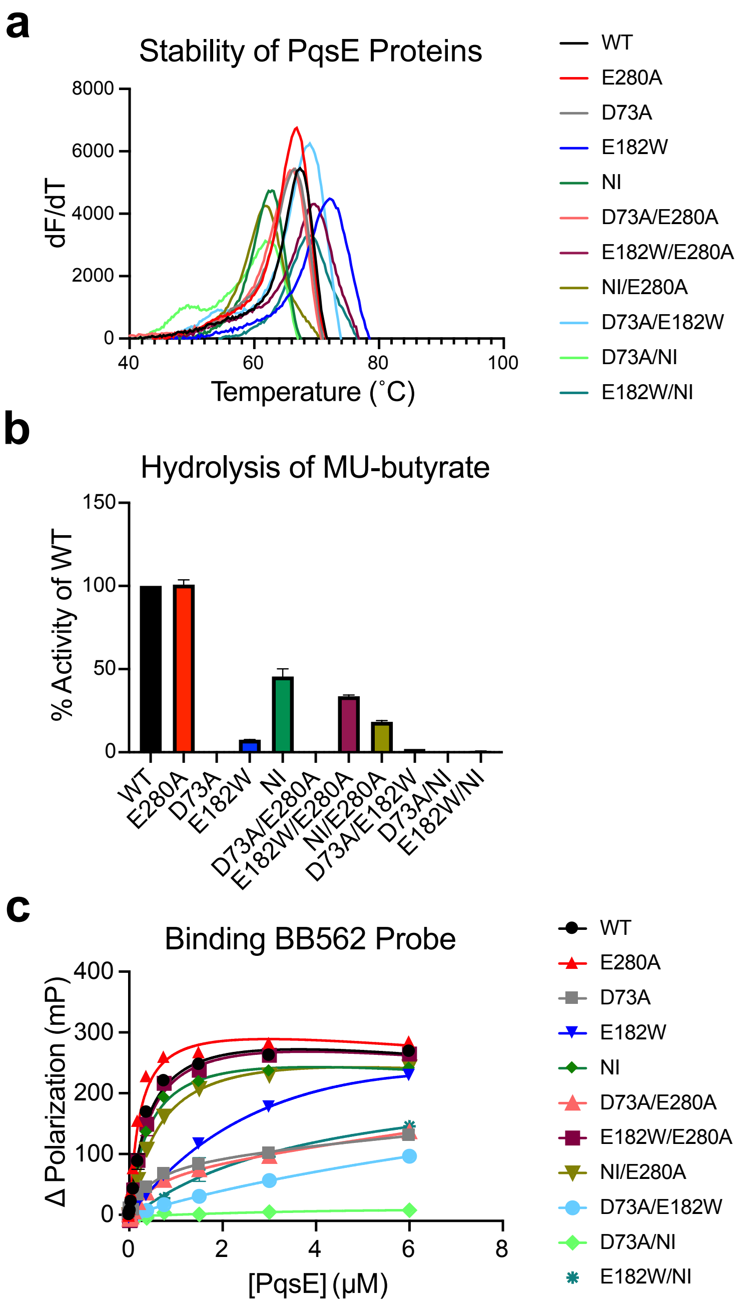

**Fig. S3.** Biochemical characterization of purified PqsE variants. a) First derivative plots of melt curves for PqsE proteins. All proteins were tested at 5 μM with 200 μM $MnCl_2$. dF/dT indicates the change in SYPRO Orange fluorescence divided by the change in temperature. The peak of each curve is interpreted as the $T_m$. Each melt curve was performed in triplicate and the first derivative was taken for the resulting averaged melt curves. b) Rate of hydrolysis of the substrate MU-butyrate (2 μM) was determined for each purified PqsE protein at 100 nM over 2.5 min. Results are the average of two independent experiments performed in triplicate. Rates are normalized to that of the WT PqsE protein (100%). c) Binding of PqsE proteins to the BB562 active site fluorescent probe in the presence of 2 μM $MnCl_2$ was measured as the change in fluorescence polarization. Results are the average of two independent experiments performed in triplicate. Error bars represent standard deviations.

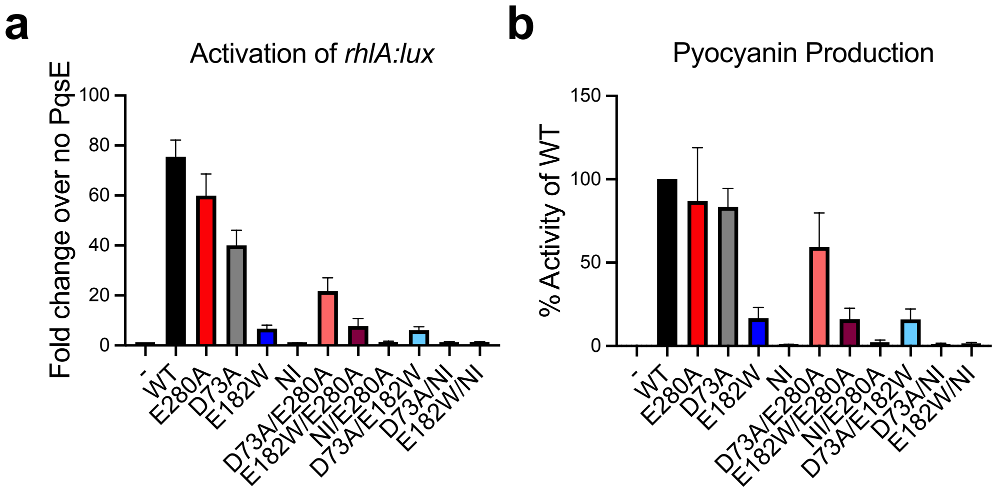

**Fig. S4.** *In vivo* characterization of PqsE proteins. a) Activation of RhlR transcription factor activity by production of RhlR and PqsE proteins in *E. coli*. All strains were grown in the presence of 100 nM C4-HSL. The bioluminescence measurements were divided by the $OD_{600}$ measurements and are represented as the fold increase compared to the no PqsE strain ("-"). Results are the average of 2 biological replicates performed in technical triplicate. b) Pyocyanin production was measured for strains expressing the *pqsE* allele from the pUCP18 plasmid. The $OD_{695}$ values of the cell-free culture fluids were measured and divided by the $OD_{600}$ values of the cultures. The results were normalized to that of the strain producing WT PqsE (100%). Results are the average of 2 biological replicates. Error bars represent standard deviations.

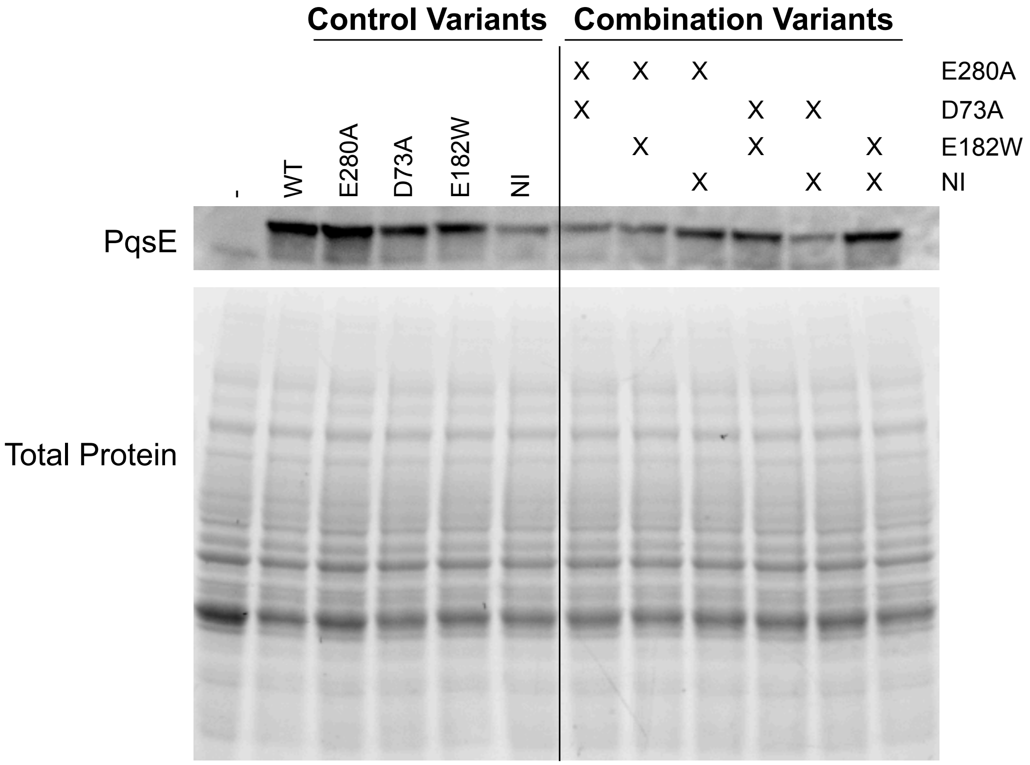

**Fig. S5.** Western blot analysis of PqsE proteins present in *E. coli* lysates. Lysates from *E. coli* strains producing the indicated PqsE protein from the pACYC184 plasmid (or harboring the empty vector, designated "-") were prepared. Top: Western blot of the gel (shown on the bottom) following incubation with a primary rabbit polyclonal $\alpha$-PqsE antibody and secondary $\alpha$-rabbit HRP-conjugated antibody. Detection was via chemiluminescence. Bottom: Total protein was imaged using a UV-activated in-gel protein stain. For the combination variants, "X" signifies that the variant in that lane harbors the corresponding substitution indicated on the right.

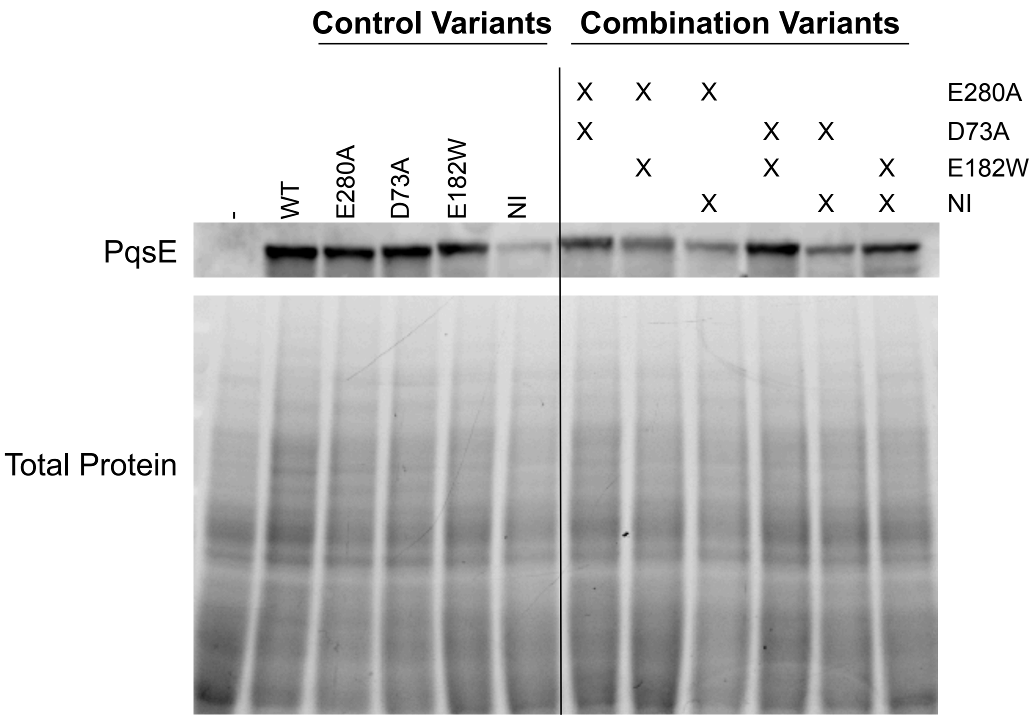

**Fig. S6.** Western blot analysis of *P. aeruginosa* PA14 lysate. Lysates were prepared from *P. aeruginosa* PA14 strains harboring *pqsE* variants on the pUCP18 plasmid (or the empty vector, designated "-"). Top: Western blot of the gel (shown in the bottom) following incubation with a primary rabbit polyclonal $\alpha$-PqsE antibody and secondary $\alpha$-rabbit HRP-conjugated antibody. Detection was via chemiluminescence. Bottom: Total protein was imaged using a UV-activated in-gel protein stain. For the combination variants, "X" signifies that the variant in that lane harbors the corresponding substitution indicated on the right.

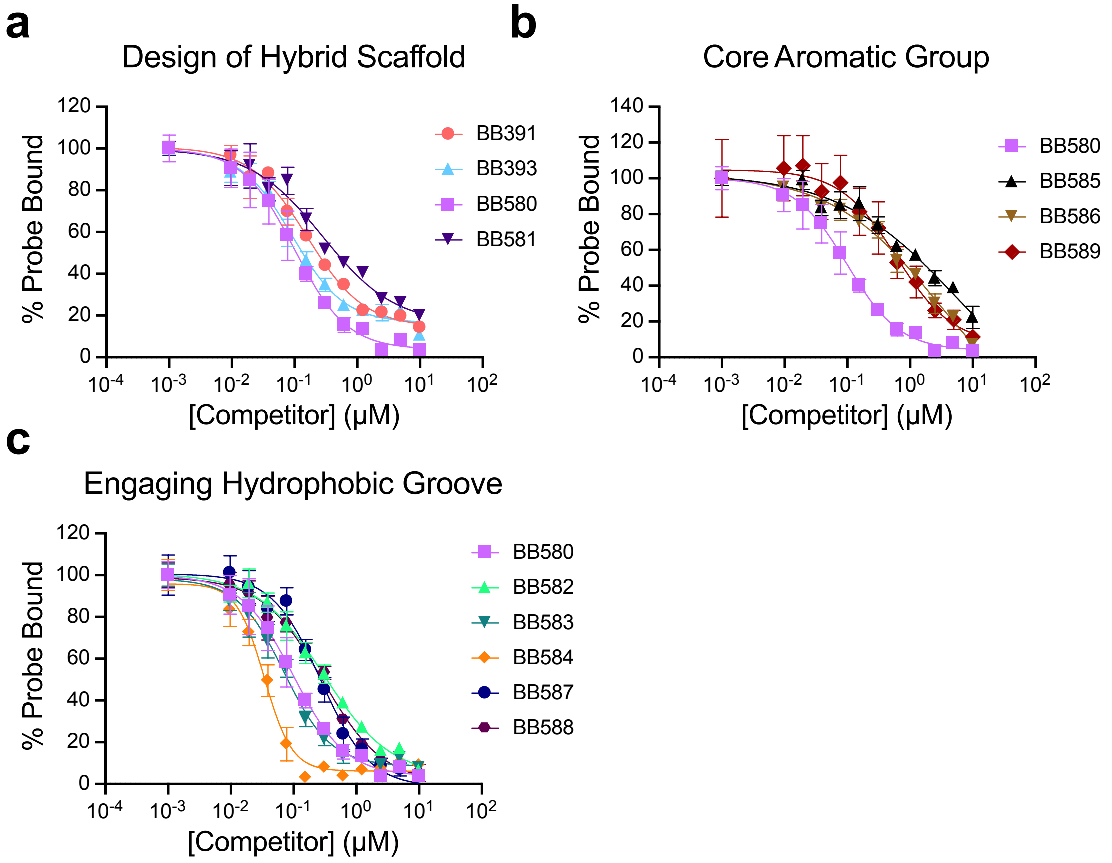

**Fig. S7.** Structure-activity relationships for BB391-BB393 hybrid derivatives. a) Optimization of “position 1”, the amide bond orientation. b) Optimization of “position 2”, the core aromatic group. c) Optimization of “position 3”, the hydrophobic tail. Competition curves with the BB562 active site fluorescent probe were fit from the average of triplicates and error bars represent standard deviations. Polarization value for PqsE-BB562 in the absence of competitor is defined as 100% Probe Bound. All polarization values were background-subtracted by the reading for the probe in the absence of PqsE (background fluorescence).

### Supplementary Tables

| Strain | Description | Reference |
|---|---|---|
| UCBPP-PA14 | PA14 *P. aeruginosa* Wildtype | Laboratory stock |
| SM52 | PA14 Δ*rhlI* | 24 |
| SM101 | PA14 Δ*rhlI* Δ*rhlR* | This study |
| SM381 | PA14 *PrhlA-mNeonGreen* | 24 |
| SM563 | PA14 Δ*pqsE PrhlA-mNeonGreen* | 25 |
| SM776 | *E. coli* BL21 (DE3) pET28b-6xHis-pqsE(WT) | 25 |
| SM1328 | *E. coli* Top10 $P_{BAD}$*-rhlR PrhlA-luxCDABE pACYC184(control)* | 20 |
| JPS0103 | *E. coli* BL21 (DE3) pET23-rhlR | 27 |
| JPS0237 | *E. coli* BL21 (DE3) pET28b-6xHis-pqsE(NI) | 21 |
| IT42 | PA14 Δ*pqsE PrhlA-mNeonGreen pUCP18-*$P_{lac}$*-pqsE(WT)* | 20 |
| IT53 | *E. coli* BL21 (DE3) pET28b-6xHis-pqsE(E182W) | 20 |
| IT68 | PA14 Δ*pqsE PrhlA-mNeonGreen pUCP18(control)* | 20 |
| IT69 | PA14 Δ*pqsE PrhlA-mNeonGreen pUCP18-*$P_{lac}$*-pqsE(E182W)* | 20 |
| IT88 | *E. coli* BL21 (DE3) pET28b-6xHis-pqsE(D73A) | 20 |
| IT89 | PA14 Δ*pqsE PrhlA-mNeonGreen pUCP18-*$P_{lac}$*-pqsE(D73A)* | 20 |
| IT100 | PA14 *pqsE(E182W)* | This study |
| IT105 | PA14 Δ*pqsE* | This study |
| IT106 | PA14 *pqsE(D73A)* | This study |
| IT107 | *E. coli* BL21 (DE3) pET28b-6xHis-pqsE(D73A/E182W) | This study |
| IT110 | *E. coli* BL21 (DE3) pET28b-6xHis-pqsE(E182W/E280A) | This study |
| IT111 | PA14 *pqsE(NI)* | This study |
| IT114 | *E. coli* BL21 (DE3) pET28b-6xHis-pqsE(E280A) | This study |
| IT116 | *E. coli* Top10 $P_{BAD}$*-rhlR PrhlA-luxCDABE pACYC184_DP-*$P_{lac}$*-pqsE(WT)* | 21 |
| IT117 | *E. coli* Top10 $P_{BAD}$*-rhlR PrhlA-luxCDABE pACYC184_DP-*$P_{lac}$*-pqsE(D73A)* | 21 |
| IT118 | *E. coli* Top10 $P_{BAD}$*-rhlR PrhlA-luxCDABE pACYC184_DP-*$P_{lac}$*-pqsE(E182W)* | 21 |
| IT119 | *E. coli* Top10 $P_{BAD}$*-rhlR PrhlA-luxCDABE pACYC184_DP-*$P_{lac}$*-pqsE(NI)* | 21 |
| IT120 | *E. coli* Top10 $P_{BAD}$*-rhlR PrhlA-luxCDABE pACYC184_DP-*$P_{lac}$*-pqsE(D73A/E182W)* | This study |
| IT121 | *E. coli* Top10 $P_{BAD}$*-rhlR PrhlA-luxCDABE pACYC184_DP-*$P_{lac}$*-pqsE(E182W/E280A)* | This study |
| IT122 | PA14 Δ*rhlI* ΔpqsE | This study |
| IT123 | PA14 Δ*rhlI pqsE(D73A)* | This study |
| IT124 | PA14 Δ*rhlI pqsE(E182W)* | This study |
| IT129 | PA14 Δ*rhlI pqsE(NI)* | This study |
| IT130 | PA14 Δ*pqsE PrhlA-mNeonGreen pUCP18-*$P_{lac}$*-pqsE(E280A)* | This study |
| IT131 | PA14 Δ*pqsE PrhlA-mNeonGreen pUCP18-*$P_{lac}$*-pqsE(D73A/E182W)* | This study |
| IT132 | PA14 Δ*pqsE PrhlA-mNeonGreen pUCP18-*$P_{lac}$*-pqsE(E182W/E280A)* | This study |
| IT133 | PA14 Δ*pqsE PrhlA-mNeonGreen pUCP18-*$P_{lac}$*-pqsE(NI)* | This study |
| IT139 | *E. coli* BL21 (DE3) pET28b-6xHis-pqsE(D73A/E280A) | This study |
| IT140 | *E. coli* BL21 (DE3) pET28b-6xHis-pqsE(NI/E280A) | This study |
| IT141 | PA14 Δ*pqsE PrhlA-mNeonGreen pUCP18-*$P_{lac}$*-pqsE(D73A/E280A)* | This study |
| IT142 | PA14 Δ*pqsE PrhlA-mNeonGreen pUCP18-*$P_{lac}$*-pqsE(NI/E280A)* | This study |
| IT143 | *E. coli* Top10 $P_{BAD}$*-rhlR PrhlA-luxCDABE pACYC184_DP-*$P_{lac}$*-pqsE(E280A)* | This study |
| IT160 | *E. coli* BL21 (DE3) pET28b-6xHis-pqsE(D73A/NI) | This study |
| IT161 | *E. coli* BL21 (DE3) pET28b-6xHis-pqsE(E182W/NI) | This study |
| IT164 | PA14 Δ*pqsE PrhlA-mNeonGreen pUCP18-*$P_{lac}$*-pqsE(D73A/NI)* | This study |
| IT165 | PA14 Δ*pqsE PrhlA-mNeonGreen pUCP18-*$P_{lac}$*-pqsE(E182W/NI)* | This study |
| IT167 | *E. coli* Top10 $P_{BAD}$*-rhlR PrhlA-luxCDABE pACYC184_DP-*$P_{lac}$*-pqsE(D73A/NI)* | This study |
| IT168 | *E. coli* Top10 $P_{BAD}$*-rhlR PrhlA-luxCDABE pACYC184_DP-*$P_{lac}$*-pqsE(E182W/NI)* | This study |

**Table S1.** Strains used in this study.

| | E182W | E182W-E280A | BB584 |
|---|---|---|---|
| **PDB accession** | 7TZ9 | 7U6G | 7TZA |
| **Data collection** | | | |
| **Space group** | *P3₂21* | *P3₂21* | *P3₂21* |
| **Cell dimensions** | | | |
| **a, b, c (Å)** | 60.7, 60.7, 146.10 | 60.6, 60.6, 146.0 | 61.1, 61.1, 146.3 |
| **α, β, γ (°)** | 90., 90., 120. | 90., 90., 120 | 90., 90., 120 |
| **Resolution** | 29 – 2.01 | 30 – 2.44 | 13 – 2.10 |
| **Outer shell** | 2.06 – 2.01 | 2.51 – 2.44 | 2.16 – 2.10 |
| **Observations** | 288714 | 116855 | 95747 |
| **Unique reflections** | 21945 | 12028 | 18595 |
| **Completeness (%)** | 99.8 (97.2) | 99.2 (89.9) | 97.0 (85.3) |
| **Multiplicity** | 13.2 (13.2) | 9.7 (7.8) | 5.1 (2.8) |
| **$R_{merge}$** | 0.079 (1.056) | 0.094 (0.936) | 0.075 (0.432) |
| **$R_{meas}$** | 0.083 (1.100) | 0.100 (1.000) | 0.083 (0.518) |
| **<I/σ(I)>** | 19.0 (2.7) | 14.7 (1.8) | 14.7 (2.6) |
| **$CC_{1/2}$** | 0.998 (0.972) | 0.999 (0.659) | 0.998 (0.769) |
| **Refinement** | | | |
| **Resolution (Å)** | 28 – 2.01 | 30 – 2.44 | 13 – 2.10 |
| **$R_{work}/R_{free}$** | 0.169/0.216 | 0.173/0.241 | 0.186/0.220 |
| **No. atoms** | | | |
| **Protein** | 2412 | 2332 | 2390 |
| **Ligand/ion** | 2 | 2 | 33 |
| **Water** | 109 | 34 | 103 |
| **Average B-factor (Å$^2$)** | 44.8 | 80.5 | 28.0 |
| **RMSD** | | | |
| **Bond lengths (Å)** | 0.006 | 0.007 | 0.007 |
| **Bond angles (°)** | 0.787 | 0.911 | 0.915 |
| **Ramachandran** | | | |
| **Favored (%)** | 96.9 | 96.5 | 97.3 |
| **Outlier (%)** | 0.0 | 0.0 | 0.0 |

**Table S2.** Crystallographic data collection and refinement statistics. Values in parentheses correspond to those for the outer shell resolution.

**Supplementary Information Text**
**Synthetic Methods**

### Abbreviations

As specified below, the symbols and conventions used in these processes, schemes, and examples are consistent with those used in the contemporary scientific literature, for example, the Journal of the American Chemical Society or the Journal of Biological Chemistry. Specifically, the following abbreviations may be used in the examples:

ACN (acetonitrile)
Boc (*tert*-butoxycarbonyl)
$CD_3OD$ (deuterated methanol)
$CDCl_3$ (deuterated chloroform)
DCM (dichloromethane)
DIEA (*N,N*-diisopropylethylamine)
DIPEA (*N,N*-diisopropylethylamine)
DMA (*N,N*-dimethylacetamide)
DMAP (4-dimethylaminopyridine)
DMF (*N,N*-dimethylformamide)
DMSO (dimethylsulfoxide)
DMSO-$d_6$ (deuterated dimethylsulfoxide)
$Et_3N$ (triethylamine)
EtOAc (ethyl acetate)
EtOH (ethanol)
FA (formic acid)
g (grams)
h (hours)
$H_2$ (hydrogen gas)
HCl (hydrochloric acid)
HOBt (hydroxybenzotriazole)
L (liters)
$LiAlH_4$ (lithium aluminum hydride)
LiOH (lithium hydroxide)
M (molar)
MeOH (methanol)
mg (milligrams)
$MgSO_4$ (magnesium sulfate)
MHz (megahertz)
min (minutes)
mL (milliliters)
mm (millimeter)
mM (millimolar)
mmol (millimoles)
mol (moles)
MTBE (methyl *tert*-butyl ether)
$N_2$ (nitrogen gas)
$Na_2SO_4$ (sodium sulfate)
Pd/C (palladium on carbon)
$Pd(OH)_2$ (palladium hydroxide)
PE (petroleum ether)
psi (pounds per square inch)
RT (room temperature)
t-Bu (*tert*-butyl)
TEA (triethylamine)
TFA (trifluoroacetic acid)
THF (tetrahydrofuran)
µL (microliters)
µm (micron)
µmol (micromoles)
BOP [(Benzotriazol-1-yloxy)tris(dimethylamino)phosphonium hexafluorophosphate]
EDCI (*N*-(3-Dimethylaminopropyl)-*N'*-ethylcarbodiimide hydrochloride)
HATU (1-[bis(dimethylamino)methylene]-1*H*-1,2,3-triazolo[4,5-b]pyridinium 3-oxide hexafluorophosphate)

### General Information

Unless otherwise indicated, all temperatures are expressed in °C (degrees Celsius). All reactions were conducted at room temperature (RT) unless otherwise noted. $^{1}$H NMR spectra were recorded on a Varian VXR-400 or a Varian Unity-400 at 400 MHz field strength. Chemical shifts are expressed in parts per million (ppm, d units). Coupling constants (*J*) are in units of Hertz (Hz). Splitting patterns describe apparent multiplicities and are designated as s (singlet), d (doublet), t (triplet), q (quartet), m (multiplet), quin (quintet), or br (broad). Mass spectral analyses were performed on a Sciex API 100 instrument using electrospray ionization (ESI). The column used for chromatography was a Luna-C18 2.0*30mm, (3 µm particles). Detection methods are diode array (DAD). MS mode was positive electrospray ionization. MS range was 100-1000. Mobile phase A was 0.037% trifluoroacetic acid in water, and mobile phase B was 0.018% trifluoroacetic acid in HPLC grade acetonitrile (10-90%) over 2 min. Analytical thin layer chromatography was used to verify the purity as well as to follow the progress of reaction(s). Unless otherwise indicated, all final products were at least 95% pure as judged by HPLC/MS.

***N*-(4-(3-(2-methyl-2-morpholinobutyl)ureido)phenyl)-1*H*-indazole-7-carboxamide (BB580)**

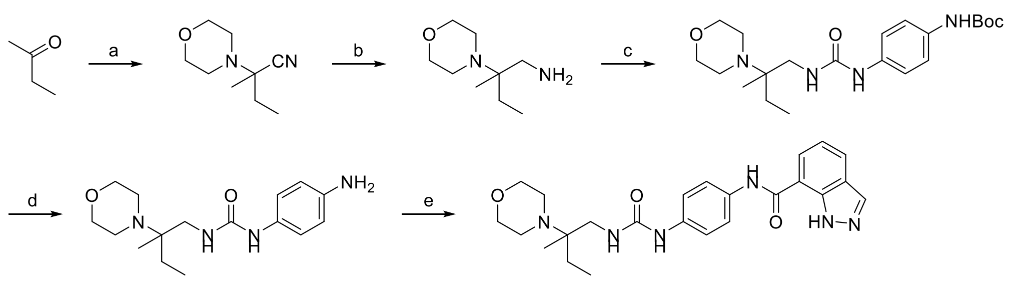

Reagents and conditions: (a) morpholine, 2-hydroxy-2-methylpropanenitrile, $MgSO_4$, DMA, 45 °C, 12 h, 28% yield; (b) $LiAlH_4$, THF, 0 °C, 12 h, 54% yield; (c) triphosgene, *tert*-butyl (4-aminophenyl)carbamate, TEA, DCM, 0 °C, 2 h, crude; (d) HCl/MeOH, RT, 1 h, 6% yield for 2 steps; (e) 1*H*-indazole-7-carboxylic acid, DIEA, HATU, DMF, RT, 1 h, 30% yield for 2 steps.

Step a: 2-methyl-2-morpholinobutanenitrile

To a solution of morpholine (9.1 g, 104 mmol, 1.5 eq), butan-2-one (5.0 g, 69.34 mmol, 1.0 eq), and 2-hydroxy-2-methylpropanenitrile (5.9 g, 69.34 mmol, 1.0 eq) in DMA (10 mL) at RT under $N_2$(g) was added $MgSO_4$ (16.7 g, 138.69 mmol, 2.0 eq). The resulting reaction mixture was stirred at 45 °C for 12 h. The reaction mixture was poured into $H_2O$ (50 mL) and extracted with EtOAc (3 x 50 mL). The combined organic phase was dried over anhydrous $Na_2SO_4$ and filtered, and the filtrate was concentrated under reduced pressure. The residue was purified by silica gel column chromatography (PE:EtOAc = 10:1 to 3:1) to afford 2-methyl-2-morpholinobutanenitrile (3.3 g, 28% yield) as a white solid. $^1H$ NMR (400 MHz, $CDCl_3$) δ 3.72 (t, *J* = 4.8 Hz, 4H), 2.64 (t, *J* = 4.8 Hz, 4H), 1.84 (q, *J* = 7.6 Hz, 2H), 1.45 (s, 3H), 1.02 (t, *J* = 7.6 Hz, 3H); LCMS calculated for $C_9H_{16}N_2O$: m/z = 168; found: m/z = 169 $(M+H)^+$.

Step b: 2-methyl-2-morpholinobutan-1-amine

To a solution of 2-methyl-2-morpholinobutanenitrile (1.5 g, 8.92 mmol, 1.0 eq) in THF (20 mL) at 0 °C under $N_2$(g) was added $LiAlH_4$ (1.18 g, 31.12 mmol, 3.49 eq). The resulting reaction mixture was stirred at RT for 12 h. The reaction mixture was poured into $H_2O$ (50 mL) at 0 °C and extracted with EtOAc (3 x 50 mL). The combined organic phase was dried over anhydrous $Na_2SO_4$ and filtered, and the filtrate was concentrated under reduced pressure to afford 2-methyl-2-morpholinobutan-1-amine (820 mg, 54% yield) as a white solid. $^1H$ NMR (400 MHz, $CDCl_3$) δ 3.63 (m, 4H), 2.40 (m, 6H), 1.51 (m, 1H), 1.22 (m, 1H), 0.92 (d, *J* = 6.4 Hz, 3H), 0.83 (t, *J* = 7.6 Hz, 3H); LCMS calculated for $C_9H_{20}N_2O$: m/z = 172; found: m/z = 173 $(M+H)^+$.

Step c: *tert*-butyl (4-(3-(2-methyl-2-morpholinobutyl)ureido)phenyl)carbamate

To a solution of triphosgene (364 mg, 1.23 mmol, 0.33 eq) in DCM (5 mL) at 0 °C under $N_2$(g) was added a solution of *tert*-butyl (4-aminophenyl)carbamate (774 mg, 3.72 mmol, 1.0 eq) in DCM (5 mL). The resulting reaction mixture was stirred at 0 °C for 1 h. Then 2-methyl-2-morpholinobutan-1-amine (640 mg, 3.72 mmol, 1.0 eq) and TEA (752 mg, 7.43 mmol, 2.0 eq) were added, and the mixture was stirred at 0 °C for 1 h. The reaction mixture was poured into $H_2O$ (50 mL) and extracted with EtOAc (3 x 30 mL). The combined organic phase was dried over anhydrous $Na_2SO_4$ and filtered, and the filtrate was concentrated under reduced pressure. The crude product was purified by reversed-phase HPLC (120 g Agela C18, 50 mL/min, water, 45-75% 20 min; 75% 10 min, Biotage) to afford *tert*-butyl (4-(3-(2-methyl-2-morpholinobutyl)ureido)phenyl)carbamate (2 g, crude) as a yellow solid. LCMS calculated for $C_{21}H_{34}N_4O_4$: m/z = 406; found: m/z = 407 $(M+H)^+$.

Step d: 1-(4-aminophenyl)-3-(2-methyl-2-morpholinobutyl)urea

To a solution of HCl/MeOH (4 M, 100 mL, 95.65 eq) at RT under $N_2$(g) was added *tert*-butyl (4-(3-(2-methyl-

2-morpholinobutyl)ureido)phenyl)carbamate (1.7 g, 4.18 mmol, 1.0 eq). The resulting reaction mixture was stirred at RT for 1 h. The reaction mixture was filtered, and the filtrate was concentrated under reduced pressure. The crude product was purified by reverse-phase preparative HPLC (column: Phenomenex Luna C18 200*40mm*10µm; mobile phase: [water(0.2%FA)-ACN]; B%: 1%-10%, 8 min) to afford 1-(4-aminophenyl)-3-(2-methyl-2-morpholinobutyl)urea (80 mg, 6% yield) as a black solid. $^1$H NMR (400 MHz, $CD_3OD$) δ 8.26 (br, s, 1H), 7.10 (d, *J* = 8.4 Hz, 2H), 6.74 (d, *J* = 8.8 Hz, 2H), 3.81 (s, 4H), 3.01 (s, 4H), 1.68 (m, 2H), 1.18 (s, 3H), 0.99 (t, *J* = 7.6 Hz, 3H); LCMS calculated for $C_{16}H_{26}N_4O_2$: m/z = 306; found: m/z = 307 $(M+H)^+$.

Step e: *N*-(4-(3-(2-methyl-2-morpholinobutyl)ureido)phenyl)-1*H*-indazole-7-carboxamide (BB580)

To a solution of 1-(4-aminophenyl)-3-(2-methyl-2-morpholinobutyl)urea (50 mg, 163 µmol, 1.0 eq) and 1*H*-indazole-7-carboxylic acid (40 mg, 244 µmol, 1.5 eq) in DMF (1 mL) at RT under $N_2$(g) was added DIEA (53 mg, 407 µmol, 2.5 eq) and HATU (93 mg, 244 µmol, 1.5 eq). The resulting reaction mixture was stirred at RT for 1 h. The reaction mixture was filtered, and the filtrate was purified by reverse-phase preparative HPLC (column: Waters Xbridge BEH C18 100*30mm*10µm; mobile phase: [water(10mM $NH_4HCO_3$)-ACN]; B%: 20%-50%,10 min) to afford *N*-(4-(3-(2-methyl-2-morpholinobutyl)ureido)phenyl)-1*H*-indazole-7-carboxamide (33.4 mg, 30% yield, 100% purity by LCMS) as a white solid. $^1$H NMR (400 MHz, DMSO-$d_6$) δ 13.11 (br, s, 1H), 10.23 (br, s, 1H), 8.63 (s, 1H), 8.12 (s, 1H), 7.96 (m, 2H), 7.59 (d, *J* = 8.8 Hz, 2H), 7.33 (d, *J* = 8.8 Hz, 2H), 7.18 (t, *J* = 7.6 Hz, 1H), 5.86 (t, *J* = 5.2 Hz, 1H), 3.53 (t, *J* = 4.4 Hz, 4H), 3.25 (s, 6H), 3.04 (dd, *J* = 18.4, 5.2 Hz, 2H), 1.35 (m, 2H), 0.84 (s, 3H), 0.75 (t, *J* = 7.6 Hz, 3H); LCMS calculated for $C_{24}H_{30}N_6O_3$: m/z = 450; found: m/z = 451 $(M+H)^+$.

**_N_-(1_H_-indazol-7-yl)-4-(3-(2-methyl-2-morpholinobutyl)ureido)benzamide (BB581)**

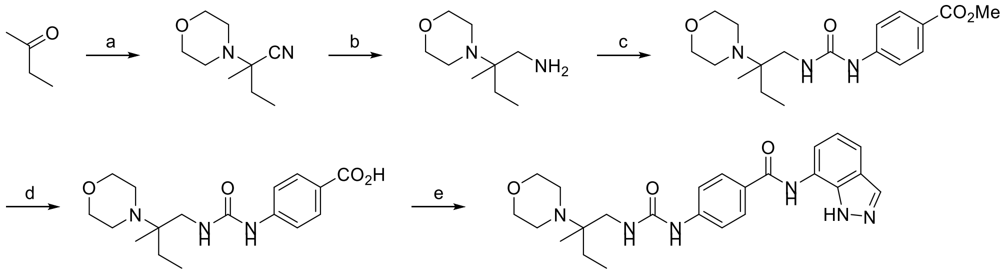

**BB0581**

Reagents and conditions: (a) morpholine, 2-hydroxy-2-methylpropanenitrile, $MgSO_4$, DMA, 45 °C, 12 h, 27% yield; (b) $LiAlH_4$, THF, 0 °C, 12 h, 26% yield; (c) methyl 4-aminobenzoate, (4-nitrophenyl) carbonochloridate, THF, RT, 12 h, 59% yield; (d) $LiOH.H_2O$, THF, 60 °C, 12 h, 30% yield; (e) 1*H*-indazol-7-amine, DIEA, HATU, DMF, 80 °C, 1 h, 37% yield.

Step a: 2-methyl-2-morpholinobutanenitrile

To a solution of morpholine (18.1 g, 208 mmol, 1.5 eq), butan-2-one (10.0 g, 138.69 mmol, 1.0 eq), and 2-hydroxy-2-methylpropanenitrile (11.8 g, 138.69 mmol, 1.0 eq) in DMA (20 mL) at RT under $N_2$(g) was added $MgSO_4$ (33.4 g, 277.37 mmol, 2.0 eq). The resulting reaction mixture was stirred at 45 °C for 12 h. The reaction mixture was poured into $H_2O$ (100 mL) and extracted with EtOAc (3 x 100 mL). The combined organic phase was dried over anhydrous $Na_2SO_4$ and filtered, and the filtrate was concentrated under reduced pressure. The residue was purified by silica gel column chromatography (PE:EtOAc = 10:1 to 3:1) to afford 2-methyl-2-morpholinobutanenitrile (6.3 g, 27% yield) as a colorless oil. $^1$H NMR (400 MHz, $CDCl_3$) δ 3.68 (t, *J* = 4.8 Hz, 4H), 2.55 (t, *J* = 4.4 Hz, 4H), 1.75 (q, *J* = 7.2 Hz, 2H), 1.36 (s, 3H), 0.93 (t, *J* = 7.6 Hz, 3H); LCMS calculated for $C_9H_{16}N_2O$: m/z = 168; found: m/z = 169 $(M+H)^+$.

Step b: 2-methyl-2-morpholinobutan-1-amine

To a solution of 2-methyl-2-morpholinobutanenitrile (3.0 g, 17.83 mmol, 1.0 eq) in THF (40 mL) at 0 °C under $N_2$(g) was added $LiAlH_4$ (2.36 g, 62.23 mmol, 3.49 eq). The resulting reaction mixture was stirred at RT for 12 h. The reaction mixture was combined with another same scale batch reaction. Then the combined mixture was poured into $H_2O$ (50 mL) at 0 °C and extracted with EtOAc (3 x 50 mL). The combined organic phase was dried over anhydrous $Na_2SO_4$ and filtered, and the filtrate was concentrated under reduced pressure to afford 2-methyl-2-morpholinobutan-1-amine (1.6 g, 26% yield) as a white solid. $^1$H NMR (400 MHz, $CDCl_3$) δ 3.62 (m, 4H), 2.47 (m, 5H), 2.33 (m, 1H), 1.50 (m, 1H), 1.20 (m, 1H), 0.91 (d, *J* = 6.4 Hz, 3H), 0.83 (m, 3H); LCMS calculated for $C_9H_{20}N_2O$: m/z = 172; found: m/z = 173 $(M+H)^+$.

Step c: methyl 4-(3-(2-methyl-2-morpholinobutyl)ureido)benzoate

To a solution of methyl 4-aminobenzoate (219 mg, 1.45 mmol, 1.0 eq) in THF (5 mL) at RT under $N_2$(g) was added (4-nitrophenyl) carbonochloridate (292 mg, 1.45 mmol, 1.0 eq). The resulting reaction mixture was stirred at RT for 0.5 h. Then 2-methyl-2-morpholinobutan-1-amine (500 mg, 2.90 mmol, 2.0 eq) was added, and the mixture was stirred at RT for 11.5 h. The reaction mixture was concentrated under reduced pressure. The crude product was purified by silica gel column chromatography (PE:EtOAc = 10:1 to 3:1) to afford methyl 4-(3-(2-methyl-2-morpholinobutyl)ureido)benzoate (300 mg, 59%) as a yellow solid. LCMS calculated for $C_{18}H_{27}N_3O_4$: m/z = 349; found: m/z = 350 $(M+H)^+$.

Step d: 4-(3-(2-methyl-2-morpholinobutyl)ureido)benzoic acid

To a solution of methyl 4-(3-(2-methyl-2-morpholinobutyl)ureido)benzoate (400 mg, 1.14 mmol, 1.0 eq) in THF (32 mL) and $H_2O$ (8 mL) at RT under $N_2$(g) was added $LiOH.H_2O$ (480 mg, 11.45 mmol, 10.0 eq). The resulting reaction mixture was stirred at 60 °C for 12 h. The reaction mixture was filtered, and the filtrate was concentrated under reduced pressure. The crude product was purified by reverse-phase preparative HPLC (column: Phenomenex luna C18 250*50mm*10µm; mobile phase: [water(0.04%HCl)-ACN]; B%: 1%-30%,10 min) to afford 4-(3-(2-methyl-2-morpholinobutyl)ureido)benzoic acid (120 mg, 30% yield) as a white solid. $^1$H NMR (400 MHz, DMSO-$d_6$) δ 12.58 (br, s, 1H), 9.55 (m, 2H), 7.85 (d, *J* = 8.8 Hz, 2H), 7.52 (d, *J* = 8.8 Hz, 2H), 7.17 (m, 1H), 3.93 (m, 4H), 3.60 (m, 5H), 3.54 (m, 1H), 3.21 (m, 2H), 1.80 (m, 2H), 1.29 (s, 3H), 0.99 (t, *J* = 7.2 Hz, 3H); LCMS calculated for $C_{17}H_{25}N_3O_4$: m/z = 335; found: m/z = 336 $(M+H)^+$.

Step e: *N*-(1*H*-indazol-7-yl)-4-(3-(2-methyl-2-morpholinobutyl)ureido)benzamide (BB581)

To a solution of 4-(3-(2-methyl-2-morpholinobutyl)ureido)benzoic acid (50 mg, 149 µmol, 1.0 eq) and 1*H*-indazol-7-amine (30 mg, 223 µmol, 1.5 eq) in DMF (1 mL) at RT under $N_2$(g) was added DIEA (48 mg, 373 µmol, 2.5 eq) and HATU (85 mg, 223 µmol, 1.5 eq). The resulting reaction mixture was stirred at 80 °C for 12 h. The reaction mixture was combined with another same scale batch reaction. Then the combined mixture was filtered, and the filtrate was purified by reverse-phase preparative HPLC (column: Waters Xbridge BEH C18 100*30mm*10µm; mobile phase: [water(10mM $NH_4HCO_3$)-ACN]; B%: 15%-45%,10 min) to afford *N*-(1*H*-indazol-7-yl)-4-(3-(2-methyl-2-morpholinobutyl)ureido)benzamide (50.1 mg, 37% yield, 100% purity by LCMS) as a white solid. $^1$H NMR (400 MHz, $CD_3OD$) δ 8.11 (s, 1H), 8.01 (d, *J* = 8.4 Hz, 2H), 7.68 (d, *J* = 8.0 Hz, 1H), 7.59 (d, *J* = 8.8 Hz, 2H), 7.50 (d, *J* = 7.2 Hz, 1H), 7.19 (t, *J* = 7.6 Hz, 1H), 3.74 (t, *J* = 4.4 Hz, 4H), 3.26 (s, 2H), 2.65 (m, 4H), 1.56 (m, 2H), 1.06 (s, 3H), 0.95 (t, *J* = 7.6 Hz, 3H); LCMS calculated for $C_{24}H_{30}N_6O_3$: m/z = 450; found: m/z = 451 $(M+H)^+$.

***N*-(4-(3-(pyrazin-2-ylmethyl)ureido)phenyl)-1*H*-indazole-7-carboxamide (BB582)**

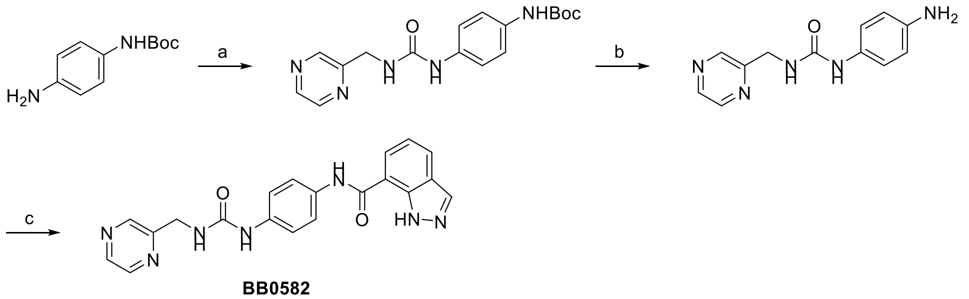

**BB0582**

Reagents and conditions: (a) triphosgene, pyrazin-2-ylmethanamine, TEA, DCM, °0 C, 2 h, 40% yield; (b) HCl/MeOH, RT, 1 h, crude; (c) 1*H*-indazole-7-carboxylic acid, DIEA, HATU, DMF, RT, 1 h, 12% yield for 2 steps.

Step a: *tert*-butyl (4-(3-(pyrazin-2-ylmethyl)ureido)phenyl)carbamate

To a solution of triphosgene (940 mg, 3.17 mmol, 0.3 eq) in DCM (10 mL) at 0 °C under $N_2$(g) was added a solution of *tert*-butyl *N*-(4-aminophenyl)carbamate (2.0 g, 9.60 mmol, 1.0 eq) in DCM (20 mL). The resulting reaction mixture was stirred at 0 °C for 1 h. Then pyrazin-2-ylmethanamine (1.05 g, 9.60 mmol, 1.0 eq) and TEA (1.94 g, 19.21 mmol, 2.0 eq) were added, and the reaction mixture was stirred at 0 °C for 1 h. The reaction mixture was poured into $H_2O$ (10 mL) and extracted with EtOAc/THF (1:1, 3 x 100 mL). The combined organic phase was dried over anhydrous $Na_2SO_4$ and filtered, and the filtrate was concentrated under reduced pressure. The crude product was triturated with EtOAc (20 mL) at RT for 10 min. The resulting slurry was filtered and the solid was collected and dried under reduced pressure to afford *tert*-butyl *N*-[4-(pyrazin-2-ylmethylcarbamoylamino)phenyl]carbamate (1.7 g, 40% yield, 78% purity by LCMS) as a black solid. $^1$H NMR (400 MHz, DMSO-$d_6$) δ 9.12 (m, 1H), 8.60 (m, 2H), 8.53 (m, 1H), 7.30 (m, 4H), 6.72 (t, *J* = 5.6 Hz, 1H), 4.45 (d, *J* = 5.6 Hz, 2H), 1.46 (m, 9H); LCMS calculated for $C_{17}H_{21}N_5O_3$: m/z = 343; found: m/z = 344 $(M+H)^+$

Step b: 1-(4-aminophenyl)-3-(pyrazin-2-ylmethyl)urea

To a solution of HCl/MeOH (4 M, 0.15 mL, 1.0 eq) at RT under $N_2$(g) was added *tert*-butyl *N*-[4-(pyrazin-2-ylmethylcarbamoylamino)phenyl]carbamate (0.2 g, 0.58 mmol, 1.0 eq). The resulting reaction mixture was stirred at RT for 1 h. The reaction mixture was concentrated under reduced pressure to afford crude 1-(4-aminophenyl)-3-(pyrazin-2-ylmethyl)urea (0.3 g, crude) as a yellow solid which was used without further purification. LCMS calculated for $C_{12}H_{13}N_5O$: m/z = 243; found: m/z = 266 $(M+Na)^+$.

Step c: *N*-(4-(3-(pyrazin-2-ylmethyl)ureido)phenyl)-1*H*-indazole-7-carboxamide (BB582)

To a mixture of 1-(4-aminophenyl)-3-(pyrazin-2-ylmethyl)urea (0.3 g, 1.23 mmol, 1.0 eq) and 1*H*-indazole-7-carboxylic acid (300 mg, 1.85 mmol, 1.5 eq) in DMF (5 mL) at RT under $N_2$(g) was added DIEA (398 mg, 3.08 mmol, 2.5 eq) and HATU (703 mg, 1.85 mmol, 1.5 eq). The resulting reaction mixture was stirred at RT for 1 h. The reaction mixture was poured into $H_2O$ (10 mL) and extracted with EtOAc (2 x 20 mL). The combined organic phase was dried over anhydrous $Na_2SO_4$ and filtered, and the filtrate was concentrated under reduced pressure to give a residue. The residue was purified by reverse-phase preparative HPLC (column: Welch Xtimate C18 100*25mm*3μm; mobile phase: [water(0.05%HCl)-ACN]; B%: 5%-35%, 8 min) to afford *N*-[4-(pyrazin-2-ylmethylcarbamoylamino)phenyl]-1*H*-indazole-7-carboxamide (61 mg, 12% yield, 96% by LCMS purity) as a yellow solid. $^1$H NMR (400 MHz, DMSO-$d_6$) δ 13.16 (s, br, 1H), 10.28 (s, br, 1H), 8.76 (s, 1H), 8.65 (s, 1H), 8.61 (s, 1H), 8.55 (s, 1H), 8.18 (s, 1H), 8.05 (d, *J* = 7.2 Hz, 1H), 8.01 (d, *J* = 8 Hz, 1H), 7.68 (d, *J* = 8.8 Hz, 2H), 7.41 (d, *J* = 8.8 Hz, 2H), 6.80 (m, 1H), 6.78 (m, 1H), 4.48 (d, *J* = 5.2 Hz, 2H); LCMS calculated for $C_{20}H_{17}N_7O_2$: m/z = 387; found: m/z = 388 $(M+H)^+$.

### ***N*-(4-(3-benzylureido)phenyl)-1*H*-indazole-7-carboxamide (BB583)**

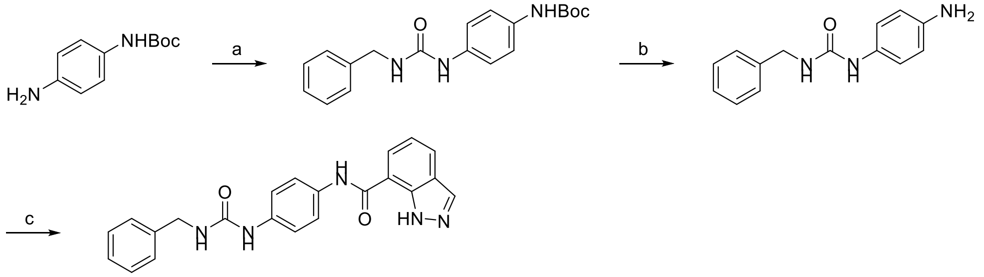

**BB0583**

Reagents and conditions: (a) triphosgene, phenylmethanamine, TEA, DCM, 0 °C, 2 h, 61% yield; (b) TFA, DCM, RT, 1 h, crude; (c) 1*H*-indazole-7-carboxylic acid, DIEA, HATU, DMF, RT, 1 h, 15% yield for 2 steps.

Step a: *tert*-butyl (4-(3-benzylureido)phenyl)carbamate

To a mixture of triphosgene (940 mg, 3.17 mmol, 0.3 eq) in DCM (10 mL) at 0 °C was added a mixture of *tert*-butyl (4-aminophenyl)carbamate (2.0 g, 9.60 mmol, 1.0 eq) in DCM (20 mL). The resulting reaction mixture was stirred for 1 h. Then a mixture of phenylmethanamine (1.03 g, 9.60 mmol, 1.0 eq) and TEA (1.94 g, 19.21 mmol, 2.0 eq) was added at 0 °C, and the reaction mixture was stirred at 0 °C for 1 h. The reaction mixture was poured into $H_2O$ (20 mL) and extracted with EtOAc/THF (1:1, 3 x 30 mL). The combined organic phase was dried over anhydrous $Na_2SO_4$ and filtered, and the filtrate was concentrated under vacuum. The residue was purified by silica gel column chromatography (PE:EtOAc = 10:1 to 0:1) to afford *tert*-butyl (4-(3-benzylureido)phenyl)carbamate (2.0 g, 61% yield, 64% purity by LCMS) as a white solid. $^1H$ NMR (400 MHz, DMSO-$d_6$) δ 9.15 (m, 1H), 8.41 (m, 1H), 7.31 (m, 9H), 6.48 (m, 1H), 4.27 (m, 2H), 1.47 (m, 9H); LCMS calculated for $C_{19}H_{23}N_3O_3$: m/z = 341; found: m/z = 342 $(M+H)^+$.

Step b: 1-(4-aminophenyl)-3-benzylurea

To a solution of *tert*-butyl (4-(3-benzylureido)phenyl)carbamate (500 mg, 1.46 mmol, 1.0 eq) in DCM (12.5 mL) at RT was added TFA (3.85 g, 33.77 mmol, 23.0 eq). The resulting reaction mixture was stirred for 1 h. The reaction mixture was concentrated under reduced pressure to afford crude 1-(4-aminophenyl)-3-benzylurea (220 mg, crude) as a yellow solid which was used without further purification. LCMS calculated for $C_{14}H_{15}N_3O$: m/z = 241; found: m/z = 242 $(M+H)^+$.

Step c: *N*-(4-(3-benzylureido)phenyl)-1*H*-indazole-7-carboxamide (BB583)

To a mixture of 1-(4-aminophenyl)-3-benzylurea (170 mg, 0.70 mmol, 1.0 eq) and 1*H*-indazole-7-carboxylic acid (171 mg, 1.06 mmol, 1.5 eq) in DMF (4 mL) at RT was added DIEA (228 mg, 1.76 mmol, 2.5 eq) and HATU (402 mg, 1.06 mmol, 1.5 eq). The resulting reaction mixture was stirred for 1 h. The reaction mixture was poured into $H_2O$ (20 mL) and extracted with EtOAc (2 x 20 mL). The combined organic phase was dried over anhydrous $Na_2SO_4$ and filtered, and the filtrate was concentrated under reduced pressure to give a residue. The residue was purified by reverse-phase preparative HPLC (column: Phenomenex Luna C18 75*30mm*3µm; mobile phase: [water(0.2%TFA)-MeOH]; B%: 40%-70%, 8 min) to afford *N*-(4-(3-benzylureido)phenyl)-1*H*-indazole-7-carboxamide (41.6 mg, 15.16% yield, 98.97% by LCMS purity) as a white solid. $^1H$ NMR (400 MHz, DMSO-$d_6$) δ 13.18 (s, br, 1H), 10.29 (s, br, 1H), 8.55 (s, 1H), 8.20 (s, 1H), 8.11 (d, *J* = 8 Hz, 1H), 8.01 (d, *J* = 8 Hz, 1H), 7.68 (d, *J* = 8.8 Hz, 2H), 7.42 (d, *J* = 8.8 Hz, 2H), 7.34 (m, 4H), 7.27 (d, *J* = 8 Hz, 2H), 6.61 (t, *J* = 6 Hz, 1H), 4.31 (d, *J* = 6 Hz, 2H); LCMS calculated for $C_{22}H_{19}N_5O_2$: m/z = 385; found: m/z = 386 $(M+H)^+$.

**_N_-(4-(3-neopentylureido)phenyl)-1_H_-indazole-7-carboxamide (BB584)**

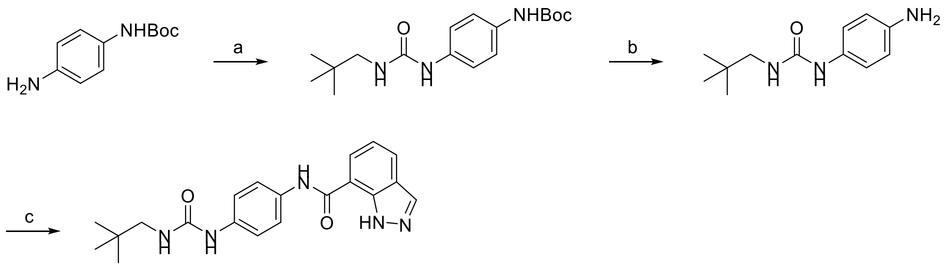

BB0584

Reagents and conditions: (a) 2,2-dimethylpropan-1-amine, triphosgene, TEA, DCM, 0 °C, 10 min; (b) TFA, DCM, RT, 1 h, crude; (c) 1*H*-indazole-7-carboxylic acid, DIEA, HATU, DMF, RT, 1 h, 45% yield for 2 steps.

Step a: *tert*-butyl (4-(3-neopentylureido)phenyl)carbamate

To a solution of triphosgene (940 mg, 3.17 mmol, 0.3 eq) in DCM (10 mL) at 0 °C under $N_2$(g) was added a solution of *tert*-butyl *N*-(4-aminophenyl)carbamate (2.0 g, 9.60 mmol, 1.0 eq) in DCM (20 mL). The resulting reaction mixture was stirred at 0 °C for 10 min. Then 2,2-dimethylpropan-1-amine (837 mg, 9.60 mmol, 1.0 eq) and TEA (1.9 g, 19.21 mmol, 2.0 eq) were added, and the mixture was stirred at 0 °C for 1 h. The reaction mixture was poured into $H_2O$ (20 mL) and extracted with EtOAc/THF (1:1, 3 x 30 mL). The combined organic phase was dried over anhydrous $Na_2SO_4$ and filtered, and the filtrate was concentrated under reduced pressure to give a residue. The residue was purified by silica gel column chromatography (PE:EtOAc = 10:1 to 0:1) to afford *tert*-butyl *N*-[4-(2,2-dimethylpropylcarbamoylamino)phenyl]carbamate (2.0 g, 65% yield) as a white solid. $^1$H NMR (400 MHz, DMSO-$d_6$) δ 9.15 (m, 1H), 8.33 (m, 1H), 7.28 (m, 4H), 5.90 (m, 1H), 2.90 (d, *J* = 6.8 Hz, 1H), 2.82 (d, *J* = 6.8 Hz, 1H), 1.47 (s, 2H), 0.84 (s, 9H); LCMS calculated for $C_{17}H_{27}N_3O_3$: m/z = 321; found: m/z = 322 $(M+H)^+$.

Step b: 1-(4-aminophenyl)-3-(2-morpholinobutyl)urea

To a solution of TFA (2.50 mL) in DCM (12.5 mL) at RT under $N_2$(g) was added *tert*-butyl *N*-[4-(2,2-dimethylpropylcarbamoylamino)phenyl]carbamate (500 mg, 1.56 mmol, 1.0 eq). The resulting reaction mixture was stirred at RT for 1 h. The reaction mixture was concentrated under reduced pressure to afford 1-(4-aminophenyl)-3-(2,2-dimethylpropyl)urea (300 mg, crude, 50% purity by LCMS) was as a yellow solid which was used without further purification. LCMS calculated for $C_{12}H_{19}N_3O$: m/z = 221; found: m/z = 222 $(M+H)^+$.

Step c: *N*-(4-(3-(2-morpholinobutyl)ureido)phenyl)-1*H*-indazole-7-carboxamide (BB584)

To a solution of 1-(4-aminophenyl)-3-(2,2-dimethylpropyl)urea (250 mg, 1.13 mmol, 1.0 eq) and 1*H*-indazole-7-carboxylic acid (274 mg, 1.69 mmol, 1.5 eq) in DMF (5 mL) at RT under $N_2$(g) was added DIEA (365 mg, 2.82 mmol, 2.5 eq) and HATU (644 mg, 1.69 mmol, 1.5 eq). The resulting reaction mixture was stirred at RT for 1 h. The reaction mixture was poured into $H_2O$ (20 mL) and extracted with EtOAc (2 x 20 mL). The combined organic phase was dried over anhydrous $Na_2SO_4$, filtered, and the filtrate was concentrated under reduced pressure to give a residue. The residue was purified by reverse-phase preparative HPLC (column: Phenomenex Gemini-NX 150*30mm*5μm; mobile phase: [water(0.2%FA)-ACN]; B%: 30%-65%, 8 min) to afford *N*-[4-(2,2-dimethylpropylcarbamoylamino)phenyl]-1*H*-indazole-7-carboxamide (187.7 mg, 45% yield, 99.88% purity by LCMS) as a white solid. $^1$H NMR (400 MHz, $CD_3OD$) δ 13.17 (s, br, 1H), 10.26 (s, br, 1H), 8.36 (s, 1H), 8.16 (s, 1H), 8.00 (m, 2H), 7.65 (d, *J* = 8.8 Hz, 2H), 7.37 (d, *J* = 8.8 Hz, 2H), 7.24 (d, *J* = 7.6 Hz, 1H), 6.13 (t, *J* = 6.4 Hz, 1H), 2.90 (d, *J* = 6, 2H), 0.85 (s, 9H); LCMS calculated for $C_{20}H_{23}N_5O_2$: m/z = 365; found: m/z = 366 $(M+H)^+$.

***N*-[5-[(2-methyl-2-morpholino-butyl)carbamoylamino]thiazol-2-yl]-1*H*-indazole-7-carboxamide (BB585)**

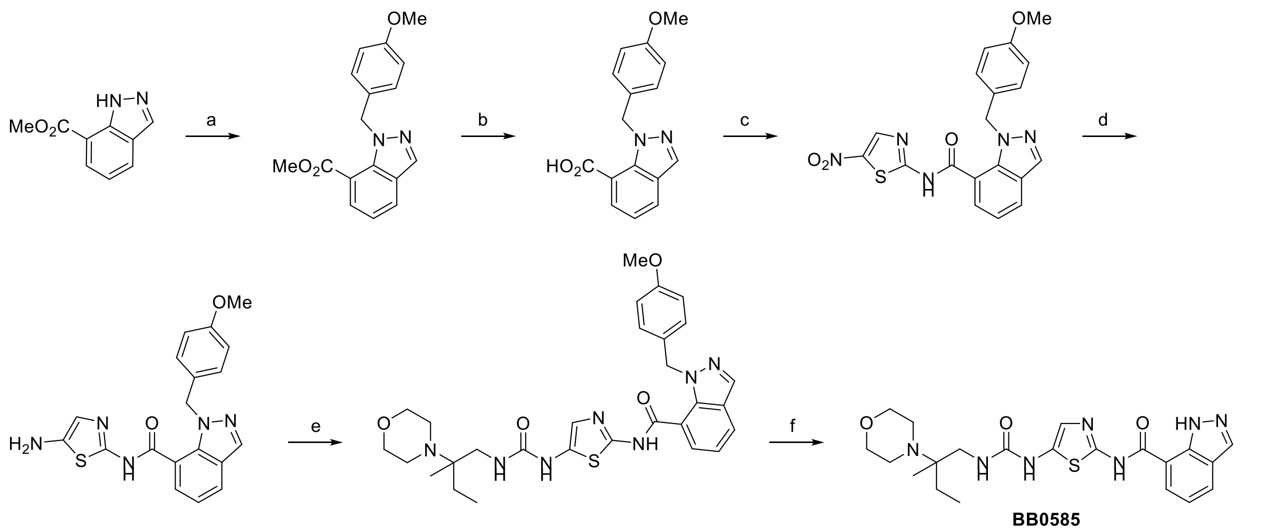

Reagents and conditions: (a) NaH, PMB-Cl, DMF, 0 °C-RT,12.5 h, 65.40% yield; (b) $LiOH.H_2O$, MeOH, $H_2O$, RT, 12 h, crude; (c) 5-nitrothiazol-2-amine, BOP, DIEA, DMF, RT, 12 h, 70.72% yield; (d) Pd/C, $H_2$ (15 psi), MeOH, RT, 12 h, crude; (e) 2-methyl-2-morpholino-butan-1-amine, bis(2,5-dioxopyrrolidin-1-yl)carbonate, pyridine, MeCN, RT, 14 h, 42% yield; (f) TFA, 100 °C, 30 min, microwave, 20% yield.

Step a: Methyl 1-[(4-methoxyphenyl)methyl]indazole-7-carboxylate

To a solution of methyl 1*H*-indazole-7-carboxylate (2 g, 11.35 mmol, 1 eq) in DMF (10 mL) was added NaH (60% in mineral oil, 590.33 mg, 14.76 mmol, 1.3 eq) at 0 °C. The reaction was stirred at 0 °C for 0.5 h, then PMB-Cl (1.78 g, 11.35 mmol, 1.55 mL, 1 eq) was added. The solution was stirred at RT for 12 h, then was poured into water (60 mL) and extracted with ethyl acetate (3 x 40 mL). The combined organic phases were washed with brine (50 mL), dried over anhydrous $Na_2SO_4$, filtered, and concentrated in vacuum. The residue was purified by flash silica gel chromatography (ISCO®; 20 g SepaFlash® silica flash column, 30-100% ethyl acetate/petroleum ether gradient at 60 mL/min) to give methyl 1-[(4-methoxyphenyl)methyl]indazole-7-carboxylate (2.2 g, 7.42 mmol, 65.40% yield) as a yellow oil.

Step b: 1-[(4-methoxyphenyl)methyl]indazole-7-carboxylic acid

$LiOH.H_2O$ (404.99 mg, 9.65 mmol, 1.3 eq) was added to a solution of methyl 1-[(4-methoxyphenyl)methyl]indazole-7-carboxylate (2.2 g, 7.42 mmol, 1 eq) in MeOH (10 mL) and $H_2O$ (5 mL) at RT, then the reaction was stirred at RT for 12 h. The solution was concentrated under reduced pressure to remove the solvent. The residue was diluted with $H_2O$ (10 mL), and the pH value of the solution was adjusted to 4 with HCl (2M). Some solid precipitated, and the mixture was filtered. The filter cake was collected and dried in vacuum to give 1-[(4-methoxyphenyl)methyl]indazole-7-carboxylic acid (1.5 g, crude) as a yellow solid. LCMS calculated for $C_{16}H_{14}N_2O_3$: m/z = 282; found: m/z = 281 $(M-1)^-$. This material was used in the next step without further purification.

Step c: 1-[(4-methoxyphenyl)methyl]-*N*-(5-nitrothiazol-2-yl)indazole-7-carboxamide

To a solution of 1-[(4-methoxyphenyl)methyl]indazole-7-carboxylic acid (0.78 g, 2.76 mmol, 1 eq) in DMF (5 mL) was added 5-nitrothiazol-2-amine (481.24 mg, 3.32 mmol, 1.2 eq), BOP (1.34 g, 3.04 mmol, 1.1 eq), and DIEA (892.77 mg, 6.91 mmol, 1.20 mL, 2.5 eq). The suspension was stirred at RT for 12 h then was filtered. The filter cake was washed with MeOH (5 mL) and dried under reduced pressure to give 1-[(4-methoxyphenyl)methyl]-*N*-(5-nitrothiazol-2-yl)indazole-7-carboxamide (0.8 g, 1.95 mmol, 70.72% yield) as a yellow solid. $^1H$ NMR (400 MHz, DMSO-$d_6$) δ 13.01 (br s, 1H), 8.83 (s, 1H), 8.75 (s, 1H), 8.21 (m, 2H), 7.43 (d, *J* = 8.4 Hz, 2H), 7.34 (dd, *J* = 8.8, 7.6 Hz, 1H), 6.95 (d, *J* = 8.8 Hz, 2H), 5.75 (s, 2H), 3.73 (s, 3H). This material was used in the next step without further purification.

Step d: *N*-(5-aminothiazol-2-yl)-1-[(4-methoxyphenyl)methyl]indazole-7-carboxamide

To a solution of 1-[(4-methoxyphenyl)methyl]-*N*-(5-nitrothiazol-2-yl)indazole-7-carboxamide (1 g, 2.44 mmol, 1 eq) in MeOH (150 mL) was added 10% Pd/C (1 g) under $N_2$. The suspension was degassed under vacuum and purged with $H_2$ several times. The suspension was stirred under $H_2$ (15 psi) at RT for 12 h. The suspension was filtered, and the filtrate was concentrated under reduced pressure to give *N*-(5-aminothiazol-2-yl)-1-[(4-methoxyphenyl)methyl]indazole-7-carboxamide (0.26 g, 685.23 μmol) as a yellowish solid. The crude product was used in the next step without further purification.

Step e: 1-[(4-methoxyphenyl)methyl]-*N*-[5-[(2-methyl-2-morpholino-butyl)carbamoylamino]thiazol-2-yl]indazole-7-carboxamide

To a solution of *N*-(5-aminothiazol-2-yl)-1-[(4-methoxyphenyl)methyl]indazole-7-carboxamide (230 mg, 606.16 μmol, 1 eq) in MeCN (5 mL) was added pyridine (47.95 mg, 606.16 μmol, 48.93 μL, 1 eq) and bis(2,5-dioxopyrrolidin-1-yl) carbonate (155.28 mg, 606.16 μmol, 1 eq) in one portion under $N_2$. The solution was stirred at 20 °C for 2 h, then 2-methyl-2-morpholino-butan-1-amine (125.31 mg, 727.40 μmol, 1.2 eq) was added, and the reaction was stirred at 20 °C for 12 h. The solution was poured into water (20 mL) and extracted with ethyl acetate (2 x 15 mL). The combined organic phases were washed with brine (10 mL), dried over anhydrous $Na_2SO_4$, filtered, and concentrated in vacuum. The residue was purified by flash silica gel chromatography (ISCO®; 4 g SepaFlash® silica flash column, 0-100% ethyl acetate/petroleum ether gradient at 50 mL/min) to afford 1-[(4-methoxyphenyl)methyl]-*N*-[5-[(2-methyl-2-morpholino-butyl)carbamoylamino]thiazol-2-yl]indazole-7-carboxamide (160 mg, 46% yield) as a yellow solid. LCMS calculated for $C_{29}H_{35}N_7O_4S$: m/z = 577; found: m/z = 578 $(M+H)^+$.

Step f: *N*-[5-[(2-methyl-2-morpholino-butyl)carbamoylamino]thiazol-2-yl]-1*H*-indazole-7-carboxamide (BB585)

1-[(4-methoxyphenyl)methyl]-*N*-[5-[(2-methyl-2-morpholino-butyl)carbamoylamino]thiazol-2-l]indazole-7-carboxamide (140 mg, 242.34 μmol, 1 eq) was taken up in TFA (1.5 mL) in a microwave tube. The tube was sealed and heated at 100 °C for 30 min under microwave. The reaction solution was concentrated under reduced pressure to remove the solvent. The residue was purified by prep-HPLC (HCl condition: column: Phenomenex Luna C18 80*40mm*3μm; mobile phase: [water(HCl)-ACN]; B%: 6%-25%,7 min) to afford *N*-[5-[(2-methyl-2-morpholinobutyl)carbamoylamino]thiazol-2-yl]-1*H*-indazole-7-carboxamide (26.3 mg, 95.6% purity, HCl salt) as an off-white solid. $^1H$ NMR (400 MHz, $CD_3OD$) δ 8.28 (s, 1H), 8.19 (br d, *J* = 7.2 Hz, 1H), 8.17 (br d, *J* = 8 Hz, 1H), 7.36 (t, *J* = 8 Hz, 1H), 7.27 (s, 1H), 4.14 (m, 2H), 3.91 (br t, *J* = 12 Hz, 2H), 3.64 (m, 4H), 3.35 (br s, 1H), 3.25 (s, 1H), 1.87 (m, 2H), 1.40 (s, 3H), 1.09 (t, *J* = 7.2 Hz, 3H); $^1H$ NMR (400 MHz, DMSO-$d_6$) δ 13.18 (m, 1H), 9.78 (s, 1H), 9.41 (m, 1H), 8.25 (s, 2H), 8.05 (d, *J* = 7.6 Hz, 1H), 7.25 (t, *J* = 7.6 Hz, 1H), 7.09 (s, 1H), 6.99 (m, 1H), 4.01 (m, 2H), 3.87 (m, 2H), 3.59 (m, 4H), 3.22 (m, 2H), 1.78 (m, 2H), 1.28 (s, 3H), 0.98 (t, *J* = 7.6 Hz, 3H); LCMS calculated for $C_{21}H_{27}N_7O_3S$: m/z = 457; found: m/z = 458 $(M+H)^+$.

**N-[5-[(2-methyl-2-morpholino-butyl)carbamoylamino]-1*H*-pyrazol-3-yl]-1*H*-indazole-7-carboxamide (BB586)**

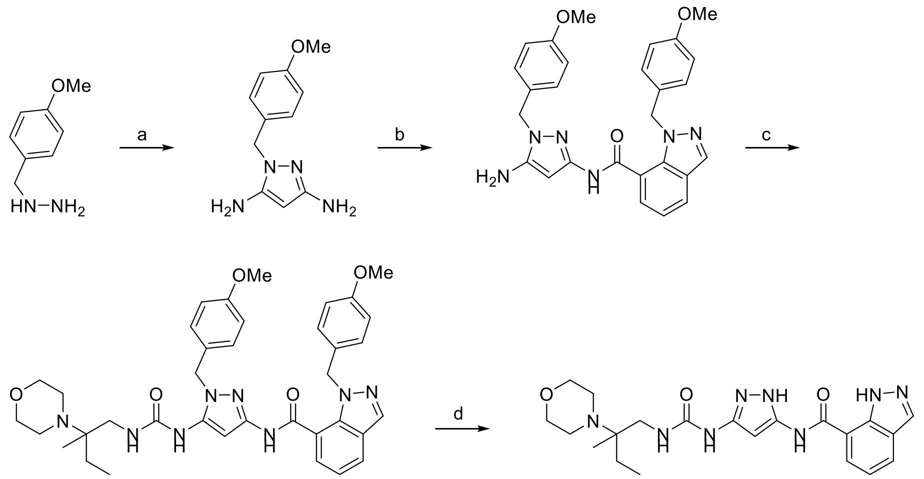

Reagents and conditions: (a) propanedinitrile, EtOH, TEA, 80 °C, 12 h, crude; (b) 1-[(4-methoxyphenyl) methyl]indazole-7-carboxylic acid, HATU, TEA, DMF, RT, 12 h, 22.52% yield; (c) 2-methyl-2-morpholino-butan-1-amine, bis(2,5-dioxopyrrolidin-1-yl)carbonate, pyridine, MeCN, RT,14 h, 19.49%; (d) TFA, 85 °C, 2 h, 11.93% yield

Step a: 1-[(4-methoxyphenyl)methyl]pyrazole-3,5-diamine

To a solution of (4-methoxyphenyl)methylhydrazine hydrochloride (2.86 g, 15.14 mmol, 1 eq) and propanedinitrile (1 g, 15.14 mmol, 952.38 µL, 1 eq) in EtOH (15 mL) was added TEA (1.23 g, 12.11 mmol, 1.69 mL, 0.8 eq), then the solution was stirred at 80 °C for 12 h under $N_2$ atmosphere. The solution was concentrated under reduced pressure to remove solvent. The residue was diluted with water (100 mL) and extracted with ethyl acetate (2 x 50 mL). The combined organic layers were washed with brine (100 mL), dried over $Na_2SO_4$, filtered, and concentrated under reduced pressure to give a residue. The residue was purified by flash silica gel chromatography (ISCO®; 20 g SepaFlash® silica flash column, 0-100% ethyl acetate/petroleum ether gradient at 80 mL/min) to afford 1-[(4-methoxyphenyl)methyl]pyrazole-3,5-diamine (3 g, crude) as brown oil. This material was used in the next step without further purification.

Step b: *N*-[5-amino-2-[(4-methoxyphenyl)methyl]pyrazol-3-yl]-1-[(4-methoxyphenyl)methyl]indazole-7-carboxamide

A suspension of 1-[(4-methoxyphenyl)methyl]indazole-7-carboxylic acid (1 g, 3.54 mmol, 1 eq), 1-[(4-methoxyphenyl)methyl]pyrazole-3,5-diamine (850.46 mg, 3.90 mmol, 1.1 eq), HATU (1.62 g, 4.25 mmol, 1.2 eq) and TEA (1.08 g, 10.63 mmol, 1.48 mL, 3 eq) in DMF (10 mL) was degassed and purged with $N_2$ 3 times, then the suspension was stirred at RT for 12 h under $N_2$ atmosphere. The solution was poured into water (50 mL) and extracted with ethyl acetate (3 x 30 mL). The combined organic phases were washed with brine (50 mL), dried over anhydrous $Na_2SO_4$, filtered, and concentrated in vacuum. The residue was purified by flash silica gel chromatography (ISCO®; 20 g SepaFlash® silica flash column, 0-100% ethyl acetate/petroleum ether gradient at 80 mL/min) to afford *N*-[5-amino-2-[(4-methoxyphenyl)methyl]pyrazol-3-yl]-1-[(4-methoxyphenyl)methyl]indazole-7-carboxamide (700 mg, 797.87 umol, 22.52% yield, 55% purity) as a brown oil.

Step c: 1-[(4-methoxyphenyl)methyl]-*N*-[1-[(4-methoxyphenyl)methyl]-5-[(2-methyl-2-morpholino-butyl)carbamoylamino]pyrazol-3-yl]indazole-7-carboxamide

To a solution of *N*-[5-amino-1-[(4-methoxyphenyl)methyl]pyrazol-3-yl]-1-[(4-methoxyphenyl)methyl]indazole-7-carboxamide (0.4 g, 828.96 µmol, 1 eq) in MeCN (1 mL) was added pyridine (65.57 mg, 828.96 µmol, 66.91 µL, 1 eq) and bis(2,5-dioxopyrrolidin-1-yl) carbonate (212.35 mg, 828.96 µmol, 1 eq) in one portion under $N_2$. The solution was stirred at RT for 2 h, then 2-methyl-2-

morpholino-butan-1-amine (171.36 mg, 994.75 µmol, 1.2 eq) was added, and the reaction was stirred at RT for 12 h. The solution was poured into water (20 mL) and extracted with ethyl acetate (2 x 15 mL). The combined organic phases were washed with brine (10 mL), dried over anhydrous $Na_2SO_4$, filtered, and concentrated in vacuum. The residue was purified by flash silica gel chromatography (ISCO®; 4 g SepaFlash® silica flash column, 0-100% ethyl acetate/petroleum ether gradient at 50 mL/min) to afford 1-[(4-methoxyphenyl)methyl]-*N*-[1-[(4-methoxyphenyl)methyl]-5-[(2-methyl-2-morpholino-butyl)carbamoylamino]pyrazol-3-yl]indazole-7-carboxamide (110 mg, 161.58 µmol, 19.49 % yield) as a yellow solid.

Step d: *N*-[5-[(2-methyl-2-morpholino-butyl)carbamoylamino]-1*H*-pyrazol-3-yl]-1*H*-indazole-7-carboxamide (BB586)

1-[(4-Methoxyphenyl)methyl]-*N*-[1-[(4-methoxyphenyl)methyl]-5-[(2-methyl-2-morpholino-butyl)carbamoylamino]pyrazol-3-yl]indazole-7-carboxamide (200 mg, 293.77 µmol, 1 eq) was taken up in TFA (2 mL), and the solution was stirred at 85 °C for 2 h. The reaction solution was concentrated under reduced pressure to remove the solvent. The residue was purified by prep-HPLC (HCl condition: column: Phenomenex Luna C18 80*40mm*3µm; mobile phase: [water(HCl)-ACN]; B%: 10%-26%,7 min) to afford *N*-[5-[(2-methyl-2-morpholino-butyl)carbamoylamino]-1*H*-pyrazol-3-yl]-1*H*-indazole-7-carboxamide (50.0 mg, 99.16% purity, HCl) as an off-white solid. $^1$H NMR (400 MHz, DMSO-$d_6$) δ 12.98 (m, 1H), 11.28 (s, 1H), 9.83 (m, 1H), 9.46 (s, 1H), 8.23 (s, 1H), 8.15 (d, *J* = 7.2 Hz, 1H), 8.05 (d, *J* = 8 Hz, 1H), 7.47 (s, 1H), 7.26 (t, *J* =8 Hz, 1H), 6.39 (s, 1H), 3.97 (s, 3H), 3.63 (m, 2H), 3.52 (m, 2H), 3.42 (m, 1H), 3.21 (m, 2H), 1.79 (m, 2H), 1.29 (s, 3H), 0.98 (t, *J* = 7.6 Hz, 3H); LCMS calculated for $C_{21}H_{28}N_8O_3$: m/z = 440; found: m/z = 441 $(M+H)^+$.

**N-(4-(3-(2-morpholinobutyl)ureido)phenyl)-1H-indazole-7-carboxamide (BB587)**

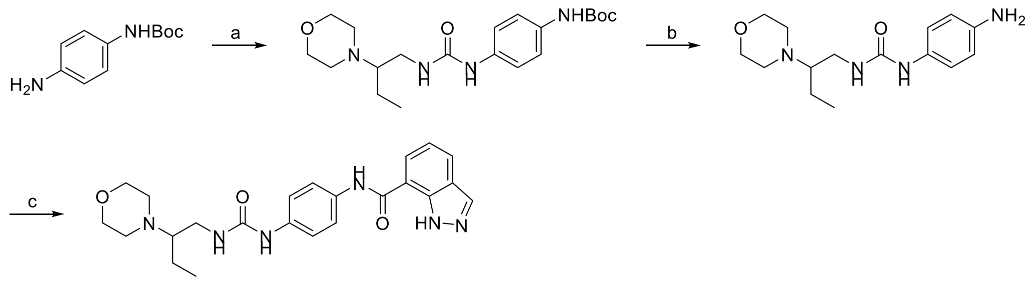

**BB0587**

Reagents and conditions: (a) 2-morpholinobutan-1-amine, triphosgene, TEA, DCM, 0 °C, 10 min, crude; (b) HCl/MeOH, RT, 1 h, crude; (c) 1*H*-indazole-7-carboxylic acid, DIEA, HATU, DMF, RT, 12 h, 5% yield for 3 steps.

Step a: *tert*-butyl (4-(3-(2-morpholinobutyl)ureido)phenyl)carbamate

To a solution of triphosgene (124 mg, 0.40 mmol, 0.3 eq) in DCM (4 mL) at 0 °C under $N_2$(g) was added a solution of *tert*-butyl *N*-(4-aminophenyl)carbamate (263 mg, 1.30 mmol, 1.0 eq) in DCM (4 mL). The resulting reaction mixture was stirred at 0 °C for 10 min. Then 2-morpholinobutan-1-amine (0.2 g, 1.26 mmol, 1.0 eq) and TEA (256 mg, 2.53 mmol, 2.0 eq) were added, and the mixture was stirred at 0 °C for 1 h. The reaction mixture was poured into $H_2O$ (10 mL) and extracted with EtOAc/THF (1:1, 3 x 20 mL). The combined organic phase was dried over anhydrous $Na_2SO_4$ and filtered, and the filtrate was concentrated under reduced pressure to afford *tert*-butyl (4-(3-(2-morpholinobutyl)ureido)phenyl)carbamate (550 mg, crude, 52% purity by LCMS) as a black solid. LCMS calculated for $C_{20}H_{32}N_4O_4$: m/z = 392; found: m/z = 393 $(M+H)^+$. This material was used without further purification.

Step b: 1-(4-aminophenyl)-3-(2-morpholinobutyl)urea

To a solution of HCl/MeOH (4 M, 15 mL) at RT under $N_2$(g) was added *tert*-butyl *N*-[4-(2 morpholinobutylcarbamoylamino)phenyl]carbamate (200 mg, 0.51 mmol, 1.0 eq). The resulting reaction mixture was stirred at RT for 1 h. The reaction mixture was concentrated under reduced pressure to afford 1-(4-aminophenyl)-3-(2-morpholinobutyl)urea (160 mg, crude, 83% purity by LCMS) as a black solid. LCMS calculated for $C_{15}H_{24}N_4O_2$: m/z = 292; found: m/z = 293 $(M+H)^+$. This material was used without further purification.

Step c: *N*-(4-(3-(2-morpholinobutyl)ureido)phenyl)-1*H*-indazole-7-carboxamide (BB587)

To a solution of 1-(4-aminophenyl)-3-(2-morpholinobutyl)urea (160 mg, 0.54 mmol, 1.0 eq) in DMF (8 mL) at RT under $N_2$(g) was added DIEA (177 mg, 1.37 mmol, 2.5 eq), HATU (312 mg, 0.82 mmol, 1.5 eq) and 1*H*-indazole-7-carboxylic acid (133 mg, 0.82 mmol, 1.5 eq). The resulting reaction mixture was stirred at RT for 16 h. The reaction mixture was concentrated under reduced pressure to give a residue. The residue was purified by reverse-phase preparative HPLC (column: Waters Xbridge BEH C18 100*30mm*10µm; mobile phase: [water(10mM $NH_4HCO_3$)-ACN]; B%: 10%-40%, 8 min) to afford *N*-(4-(3-(2-morpholinobutyl)ureido)phenyl)-1*H*-indazole-7-carboxamide (11.8 mg, 5% yield, 96% purity by LCMS) as a white solid. $^1$H NMR (400 MHz, DMSO-$d_6$) δ 13.16 (s, br, 1H), 10.27 (s, br, 1H), 8.63 (s, 1H), 8.06 (s, 1H), 8.17 (br s, 1H), 8.05 (d, *J* = 6.8 Hz, 1H), 8.01 (d, *J* = 8 Hz, 1H), 7.67 (d, *J* = 9.2 Hz, 2H), 7.40 (d, *J* = 8.8 Hz, 2H), 7.26 (t, *J* = 7.6 Hz, 1H), 6.03 (m, 1H), 3.63 (m, 4H), 3.25 (m, 1H), 3.24 (m, 1H), 2.55 (m, 3H), 2.35 (m, 2H), 1.55 (m, 1H), 1.25 (m, 1H),1.08 (t, *J* = 7.2 Hz, 3H); LCMS calculated for $C_{23}H_{28}N_6O_3$: m/z = 436; found: m/z = 437 $(M+H)^+$.

***N*-(4-(3-(2-morpholinopropyl)ureido)phenyl)-1*H*-indazole-7-carboxamide (BB588)**

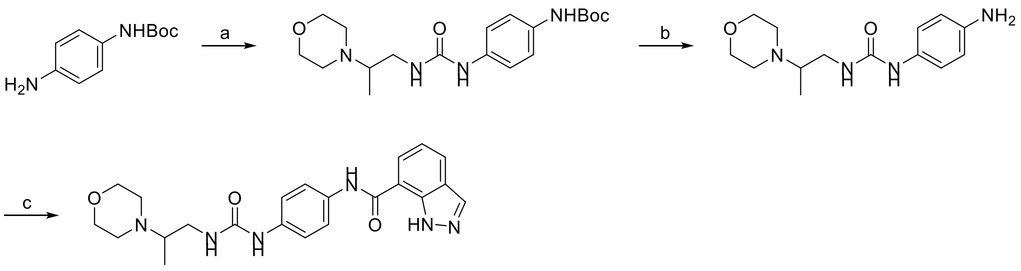

**BB0588**

Reagents and conditions: (a) 2-morpholinopropan-1-amine, triphosgene, TEA, DCM, 0 °C, 10 min, crude; (b) HCl/MeOH, RT, 1 h, crude; (c) 1*H*-indazole-7-carboxylic acid, DIEA, HATU, DMF, RT, 12 h, 30% yield for 3 steps.

Step a: *tert*-butyl (4-(3-(2-morpholinopropyl)ureido)phenyl)carbamate

To a solution of triphosgene (115 mg, 0.39 mmol, 0.3 eq) in DCM (3.5 mL) at 0 °C under $N_2$(g) was added a solution of *tert*-butyl *N*-(4-aminophenyl)carbamate (245 mg, 1.18 mmol, 1.0 eq) in DCM (3.5 mL). The resulting reaction mixture was stirred at 0 °C for 10 min. Then 2-morpholinopropan-1-amine (170 mg, 1.18 mmol, 1.0 eq) and TEA (238 mg, 2.36 mmol, 2.0 eq) were added, and the mixture was stirred at 0 °C for 1 h. The reaction mixture was poured into $H_2O$ (15 mL) and extracted with EtOAc/THF (1:1, 3 x 30 mL). The combined organic phase was dried over anhydrous $Na_2SO_4$ and filtered, and the filtrate was concentrated under reduced pressure to afford *tert*-butyl *N*-[4-(2-morpholinopropylcarbamoylamino)phenyl]carbamate (550 mg, crude, 29% purity by LCMS) as a black solid. LCMS calculated for $C_{19}H_{30}N_4O_4$: m/z = 378; found: m/z = 379 $(M+H)^+$. This material was used without further purification.

Step b: 1-(4-aminophenyl)-3-(2-morpholinopropyl)urea

To a solution of HCl/MeOH (4 M, 20 mL) at RT under $N_2$(g) was added *tert*-butyl *N*-[4-(2-morpholinopropylcarbamoylamino)phenyl]carbamate (450 mg, 1.19 mmol, 1.0 eq). The resulting reaction mixture was stirred at RT for 1 h. The reaction mixture was concentrated under reduced pressure to afford 1-(4-aminophenyl)-3-(2-morpholinopropyl)urea (330 mg, crude, 86% purity by LCMS) as a black solid.

LCMS calculated for $C_{14}H_{22}N_4O_2$: m/z = 278; found: m/z = 279 $(M+H)^+$. This material was used without further purification.

Step c: *N*-(4-(3-(2-morpholinopropyl)ureido)phenyl)-1*H*-indazole-7-carboxamide (BB588)

To a solution of 1-(4-aminophenyl)-3-(2-morpholinopropyl)urea (330 mg, 1.19 mmol, 1.0 eq) in DMF (5 mL) at RT under $N_2$(g) was added DIEA (383 mg, 2.96 mmol, 2.5 eq), HATU (676 mg, 1.78 mmol, 1.5 eq) and 1*H*-indazole-7-carboxylic acid (288 mg, 1.78 mmol, 1.5 eq). The resulting reaction mixture was stirred at RT for 12 h. The reaction mixture was concentrated under reduced pressure to give a residue. The residue was purified by reverse-phase preparative HPLC (column: Phenomenex Gemini-NX 80*40mm*3μm; mobile phase: [water(10mM $NH_4HCO_3$)-ACN]; B%: 10%-40%, 8 min) to afford *N*-[4-(2-morpholinopropylcarbamoylamino)phenyl]-1*H*-indazole-7-carboxamide (157 mg, 30% yield, 95% purity by LCMS) as a black solid. $^1$H NMR (400 MHz, $CD_3OD$) δ 8.17 (br s, 1H), 8.04 (m, 2H), 7.68 (d, *J* = 9.2 Hz, 2H), 7.41 (d, *J* = 9.2 Hz, 2H), 7.30 (t, *J* = 7.6 Hz, 1H), 3.73 (m, 4H), 3.33 (m, 1H), 3.32 (m, 1H), 2.65 (m, 3H), 2.54 (m, 2H), 1.06 (d, *J* = 6.8 Hz, 3H); LCMS calculated for $C_{22}H_{26}N_6O_3$: m/z = 422; found: m/z = 423 $(M+H)^+$.

**N-[5-[(2-methyl-2-morpholino-butyl)carbamoylamino]-2-pyridyl]-1*H*-indazole-7-carboxamide (BB589)**

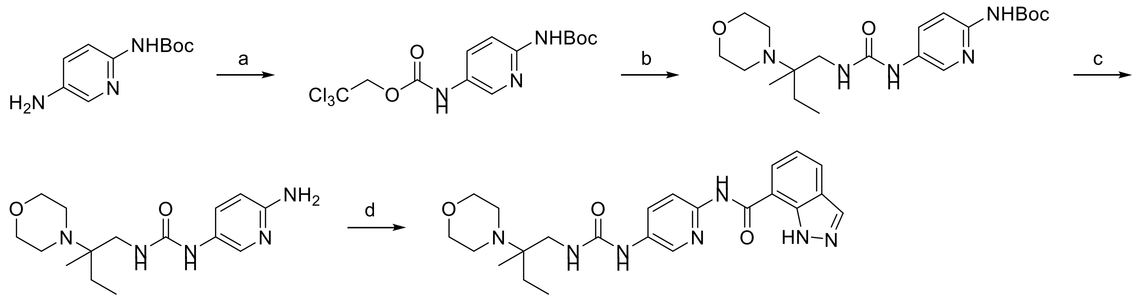

**BB0589**

Reagents and conditions: (a) 2,2,2-trichloroethyl carbonochloridate, pyridine, THF, 0 °C-RT, 2 h, crude; (b) 2-methyl-2-morpholino-butan-1-amine, DIPEA, DMSO, 70 °C, 12 h, crude; (c) HCl/EtOAc, MeOH, RT, 2 h, crude; (d) 1*H*-indazole-7-carboxylic acid, HATU, TEA, DMF, RT, 6 h, 11.93% yield.

Step a: 2,2,2-trichloroethyl *N*-[6-(*tert*-butoxycarbonylamino)-3-pyridyl]carbamate

To a solution of *tert*-butyl (5-aminopyridin-2-yl)carbamate (1 g, 4.78 mmol, 1 eq) and pyridine (45.36 mg, 573.49 μmol, 46.29 μL, 1.2 eq) in THF (20 mL) was added 2,2,2-trichloroethyl carbonochloridate (1.21 g, 5.73 mmol, 768.98 μL, 1.2 eq) at 0 °C. The suspension was stirred at RT for 2 h, then was poured into water (50 mL) and extracted with ethyl acetate (2 x 30 mL). The combined organic phases were washed with brine (50 mL), dried over anhydrous $Na_2SO_4$, filtered, and concentrated in vacuum to afford 2,2,2-trichloroethyl *N*-[6-(*tert*-butoxycarbonylamino)-3-pyridyl]carbamate (2 g, crude) as a white solid. $^1$H NMR (400 MHz, DMSO-$d_6$) δ 10.20 (br s, 1H), 9.67 (s, 1H), 8.34 (br s, 1H), 7.84 (br d, *J* = 8.4 Hz, 1H), 7.73 (d, *J* = 8.8 Hz, 1H), 4.93 (s, 2H), 1.46 (s, 9H); LCMS calculated for $C_{13}H_{16}Cl_3N_3O_4$: m/z = 384; found: m/z = 385 $(M+H)^+$. The crude product was used in the next step without further purification.

Step b: *tert*-butyl *N*-[5-[(2-methyl-2-morpholino-butyl)carbamoylamino]-2-pyridyl]carbamate

To a solution of 2,2,2-trichloroethyl *N*-[6-(*tert*-butoxycarbonylamino)-3-pyridyl]carbamate (1 g, 2.60 mmol, 1 eq) in DMSO (10 mL) was added DIPEA (436.81 mg, 3.38 mmol, 588.69 μL, 1.3 eq) and 2-methyl-2-morpholino-butan-1-amine (582.22 mg, 3.38 mmol, 1.3 eq). The solution was stirred at 70 °C for 12 h, then was poured into water (30 mL) and extracted with ethyl acetate (3 x 20 mL). The combined organic phases were washed with brine (20 mL), dried over anhydrous $Na_2SO_4$, filtered, and concentrated in vacuum. The residue was purified by flash silica gel chromatography (ISCO®; 20 g SepaFlash® silica flash column, 0-

80% ethyl acetate/petroleum ether gradient at 60 mL/min) to afford *tert*-butyl *N*-[5-[(2-methyl-2-morpholino-butyl)carbamoylamino]-2-pyridyl]carbamate (440 mg) as a yellowish solid. $^1$H NMR (400 MHz, DMSO-$d_6$) δ 9.54 (m, 1H), 8.74 (s, 1H), 8.27 (m, 1H), 7.78 (m, 1H), 7.69 (m, 1H), 5.97 (m, 1H), 3.58 (br t, *J* = 4.4 Hz, 4H), 3.42 (m, 3H), 3.32 (br s, 2H), 3.09 (m, 1H), 1.46 (m, 12H), 0.88 (m, 2H), 0.81 (t, *J* = 7.6 Hz, 2H). The crude product was used in the next step without further purification.

Step c: 1-(6-amino-3-pyridyl)-3-(2-methyl-2-morpholino-butyl)urea

To a solution of *tert*-butyl *N*-[5-[(2-methyl-2-morpholino-butyl)carbamoylamino]-2-pyridyl] carbamate (390 mg, 957.04 μmol, 1 eq) in MeOH (5 mL) was added HCl/EtOAc (10 mL). The solution was stirred at RT for 2 h, then was concentrated under reduced pressure to remove the solvent. 1-(6-amino-3-pyridyl)-3-(2-methyl-2-morpholino-butyl)urea (crude, 350 mg, HCl salt) was obtained as a brown solid. $^1$H NMR (400 MHz, DMSO-$d_6$) δ 8.20 (s, 1H), 7.89 (dd, *J* = 9.6, 2.4 Hz, 1H), 7.01 (d, *J* = 7.2 Hz, 1H), 4.09 (m, 2H), 3.89 (m, 2H), 3.64 (m, 3H), 3.35 (s, 2H), 3.27 (m, 1H), 1.85 (m, 2H), 1.37 (s, 3H), 1.07 (t, *J* = 7.6 Hz, 3H). The crude product was used in the next step without further purification.

Step d: *N*-[5-[(2-methyl-2-morpholino-butyl)carbamoylamino]-2-pyridyl]-1*H*-indazole-7-carboxamide (BB589)

To a solution of 1*H*-indazole-7-carboxylic acid (101.86 mg, 628.18 μmol, 1.2 eq) in DMF (2 mL) was added HATU (238.85 mg, 628.18 μmol, 1.2 eq) and TEA (211.88 mg, 2.09 mmol, 291.45 μL, 4 eq). The solution was stirred at 20 °C for 10 min, then 1-(6-amino-3-pyridyl)-3-(2-methyl-2-morpholino-butyl)urea (0.18 g, 523.48 μmol, 1 eq, HCl salt) was added. The solution was stirred at RT for 6 h, then was poured into water (30 mL) and extracted with ethyl acetate (2 x 20 mL). The combined organic phases were washed with brine (20 mL), dried over anhydrous $Na_2SO_4$, filtered, and concentrated in vacuum. The residue was purified by prep-HPLC (neutral condition: column: Waters Xbridge BEH C18 100*25mm*5μm; mobile phase: [water(10 mM $NH_4HCO_3$)-ACN]; B%: 20%-50%, 10 min ) to give the impure product (60 mg, purity 83% by HPLC) as a light yellow solid. The product was triturated with DMF (5 mL) at 20 °C for 10 min and filtered, and the filter cake was washed with $H_2O$ (3 mL). After lyophilization, *N*-[5-[(2-methyl-2-morpholino-butyl)carbamoylamino]-2-pyridyl]-1*H*-indazole-7-carboxamide (30.2 mg, 62.44 μmol, 11.93% yield, 93.36% purity) was obtained as a white solid. $^1$H NMR (400 MHz, $CD_3OD$) δ 8.46 (s, 1H), 8.37 (m, 1H), 8.19 (m, 3H), 7.95 (m, 1H), 7.31 (t, *J* = 7.6 Hz, 1H), 4.13 (m, 2H), 3.85 (m, 2H), 3.68 (m, 2H), 3.58 (m, 2H), 1.84 (m, 2H), 1.38 (s, 3H), 1.08 (br t, *J* = 7.2 Hz, 3H); $^1$H NMR (400 MHz, DMSO-$d_6$) δ 13.20 (br s, 1H), 10.82 (s, 1H), 9.52 (m, 1H), 9.31 (br s, 1H), 8.46 (br s, 1H), 8.20 (m, 2H), 8.02 (br d, *J* = 8 Hz, 1H), 7.92 (m, 1H), 7.19 (m, 2H), 3.95 (m, 4H), 3.56 (m, 3H), 3.42 (m, 1H), 3.20 (m, 2H), 1.79 (m, 2H), 1.28 (s, 3H), 0.98 (br t, *J* = 7.2 Hz, 3H); LCMS calculated for $C_{23}H_{29}N_7O_3$: m/z = 451; found: m/z = 452 $(M+H)^+$.
